# Meta-omic insights into active bacteria mediating N_2_O mitigation and dissimilatory nitrate reduction to ammonium in an ammonia recovery bioreactor

**DOI:** 10.1101/2024.11.13.623363

**Authors:** Hop V. Phan, Shohei Yasuda, Kohei Oba, Hiroki Tsukamoto, Tomoyuki Hori, Megumi Kuroiwa, Akihiko Terada

## Abstract

Shifting from ammonia removal to recovery is the current strategy in wastewater treatment management. We recently developed a microaerophilic activated sludge (MAS) system for retaining ammonia while removing organic carbon with minimal N_2_O emissions. A comprehensive understanding of nitrogen metabolisms in the MAS system is essential to optimize system performance. Here, we employed metagenomics and metatranscriptomics analyses to characterize the microbial community structure and activity during the transition from a microaerophilic to an aerobic condition. A hybrid approach of high-quality Illumina short reads and Nanopore long reads recovered medium-to high-quality 98 non-redundant metagenome-assembled genomes (MAGs) from the MAS communities. The suppressed bacterial ammonia monooxygenase (*amoA*) expression was upregulated after shifting from a microaerophilic to an aerobic condition. The 73 MAGs (>74% of the total) from 11 bacterial phyla harbored genes encoding proteins involved in nitrate respiration; 39 MAGs (∼53%) carried N_2_O reductase (*nosZ*) genes with the predominance of clade II *nosZ* (31 MAGs), and 24 MAGs (∼33%) possessed nitrite reductase (ammonia forming) genes (*nrfA*). Clade II *nosZ* and *nrfA* genes exhibited the highest and second-highest expressions among nitrogen metabolism genes, indicating robust N_2_O consumption and ammonification. Non-denitrifying clade II *nosZ* bacteria, *Cloacibacterium* spp., in the most abundant and active phylum Bacteroioda, were likely major N_2_O sinks. Elevated dissolved oxygen (DO) concentration inhibited clade II *nosZ* expression but not *nrfA* expression, potentially switching phenotypes from N_2_O reduction to ammonification. Collectively, the multi-omics analysis illuminated vital bacteria responsible for N_2_O reduction and ammonification in microaerophilic and aerobic conditions, facilitating high-performance ammonia recovery.

## INTRODUCTION

In conventional wastewater treatment plants, nitrogen in wastewater is removed by converting reactive nitrogen into dinitrogen gas via nitrification-denitrification. While this process alleviates the environmental burden associated with nitrogen constituents, the requirements of energy-intensive aeration, external organic carbon, and waste sludge disposal make the process incompatible with sustainable development goals [1, 2]. A transition from nitrogen removal to recovery was initiated in wastewater treatment plants toward sustainable nitrogen management. Ammonia recovery from high-strength nitrogenous organic wastewater ensures energy stored in ammonia, overcoming the high energy demand in conventional nitrogen removal and preventing the emission of nitrous oxide (N_2_O), which is a potent greenhouse gas with a global warming potential 273 times higher than that of CO_2_ and accounts for nearly 80% of the total CO_2_ footprint of a wastewater treatment plant (WWTP) [3].

We recently developed a microaerophilic activated sludge (MAS) system to retain ammonia in nitrogenous wastewater while removing organic carbon [4, 5]. Setting low dissolved oxygen (DO) concentrations (< 0.1 mg/L) and short solids retention times (SRT) (< 5 d) in a MAS system out-selects nitrifying bacteria for suppressing ammonia oxidation, which saves energy and prevents N_2_O emissions. In contrast, the fluctuation of influent nitrogen loadings increases oxygen concentrations, initiating nitrification and producing N_2_O in the system [5]. Understanding the microbial community and functions regarding carbon and nitrogen metabolisms in the MAS system is essential to optimize the system operation for removing organic carbon, enriching ammonia, and mitigating N_2_O emissions.

Mitigating N_2_O emission entails a better understanding of N_2_O-reducing bacteria. N_2_O-reducing bacteria reduce N_2_O to N_2_ anaerobically by a copper-dependent N_2_O reductase (NosZ) [6]. This enzyme is phylogenetically classified into two clades, which differ in *nos* gene clusters, translocation (clade I) or secretory (clade II) pathways [7, 8]. Clade I *nosZ* is typically present in canonical denitrifiers possessing an entire set of denitrifying genes. Over half of the bacteria containing clade II *nosZ* are non-denitrifiers, missing genes encoding nitrate and nitrite reductases [9]. Studies reported higher abundances of clade II *nosZ* bacteria in soils and WWTPs than clade I *nosZ* bacteria [7, 10], likely because of their higher competitiveness and energy efficiency under N_2_O-limited conditions. These physiological traits, however, have been controversial, claiming higher growth yields and N_2_O affinities of clade II *nosZ* bacteria [11–13] and *vice versa* [14, 15]. Thus, understanding clade I and II *nosZ* ecophysiologies is indispensable for N_2_O mitigation. Exploring active N_2_O-reducing bacteria under microaerophilic conditions in the MAS system is crucial for the sustainable mitigation of sporadically occurring N_2_O emission [5] as the MAS system employs a microaerophilic condition. Recently, multiple phenotypes of N_2_O reduction in the presence of oxygen, *i.e.*, intolerant, sensitive, and tolerant trends, were reported [12, 16–19]. Their complex phenotypes, nevertheless, are still under debate.

Bacteria exerting dissimilatory nitrate reduction to ammonia (DNRA) are likely key guilds in the MAS system for improving ammonia recovery. These bacteria possess pentaheme *c*-type (c552) cytochrome nitrite reductase (encoded by *nrfA* genes), NADH-dependent nitrite reductases (encoded by *nirB* genes), or octaheme tetrathionate reductase (encoded by *octR* genes) [20, 21]. Despite the ubiquitous presence of DNRA bacteria, studies of DNRA in WWTPs have been limited. Wang et al. [22] reported a lower abundance and activity of DNRA than denitrifying bacteria in WWTPs. Applying their functions in ammonia recovery [23, 24] requires more research.

Bacteria performing both N_2_O reduction and DNRA could be harnessed in the MAS system. Nearly one-third of non-denitrifiers with clade II *nosZ* also possess *nrfA*, suggesting the omnipresence of these specialized bacteria in the environment [8]. Pure culture works revealed the growth conditions and regulatory mechanisms between DNRA and denitrification (canonical denitrifiers) or N_2_O respiration (non-denitrifiers) [25–27]. Nevertheless, knowledge about the N_2_O-reducing and DNRA specialists, especially in engineered systems, is scarce. A comprehensive understanding of the intricate genotypes and transcripts of these specialists may allow the development of conditions to maximize N_2_O reduction and DNRA activities, enhancing the ammonia recovery performance of the MAS systems.

This study used a meta-omics approach by hybrid sequencing to elucidate the ecophysiologies of nitrogen-transforming microorganisms in a MAS system. Characterizing the microbial community structure and activity under microaerophilic and oxygen-elevating conditions is essential to manage the MAS system. We hypothesized that (i) the activity of N_2_O reducers is compromised at an elevated DO concentration, activating ammonia-oxidizing microorganisms (AOMs) to emit N_2_O; (ii) the microbial guilds responsible for N_2_O reduction contribute to the mitigation of N_2_O emission under microaerophilic conditions; and (iii) DNRA bacteria possess the metabolic potential of N_2_O reduction. To examine these hypotheses, the objectives of this study were as follows: (1) to reveal which AOMs are responsible for nitrification at an elevated DO concentration; (2) to characterize the bacteria actively reducing nitrogen oxides, particularly N_2_O under microaerobic and aerobic conditions; and (3) to unravel the phylogeny of bacteria exerting N_2_O-reducing and DNRA in the MAS system.

## MATERIALS AND METHODS

### Bioreactor operation and sampling

Two parallel MAS systems (R1 and R2) were operated for nearly 300 days. These systems were designed to retain ammonia and remove organic matter from high-strength nitrogenous wastewater by suppressing ammonia oxidation and minimizing N_2_O emissions. The operation, performance, 16S rRNA gene-based microbial profiles, and qPCR of functional genes of the systems over time were previously reported [5]. Biomass samples were collected in triplicate from R2 on day 268 under microaerophilic (control) conditions with an aeration rate of 2.0 L/min and DO of < 0.02 mg/L. The aeration rate doubled (4 L/min), concomitantly increasing DO concentration. After 19 h, triplicate biomass samples for metatranscriptomics were collected. Aeration rate, DO concentration, NH_4_, and NO_3_ concentrations were recorded every minute by a DO electrode (MultiLine^®^ Multi 3510 IDS Multi-Parameter Portable Meter, WTW, Germany) and NH_4_ /NO_3_ sensors (VARiON plus 700 IQ, WTW). Off-gas N_2_O concentration was measured by GC-MS (GCMS-QP2010 SE, Shimadzu, Kyoto, Japan). These data are presented in **Supplementary Fig. S1**.

### DNA/RNA extraction and Illumina sequencing

Fresh biomass samples were subjected to DNA extraction using a Fast DNA spin kit for soil and a FastPrep-24 instrument (MP-Biomedicals, Santa Ana, CA, USA) following the manufacturer’s protocol. DNA quality was checked by 1% agarose gel electrophoresis and a spectrophotometer (NanoDrop 2000, ThermoFisher Scientific, MA, USA). DNA concentration was determined with the Qubit dsDNA HS Assay Kit (Thermo Fisher Scientific, Waltham, MA, USA) and diluted to 50 ng/µL. Sequencing libraries were prepared with the MGIEasy FS PCR Free DNA library prep set (MGI Tech, Shenzhen, China) following the manufacturer’s protocol. The 150-bp paired-end sequencing was performed using DNBSEQ-G400RS (MGI Tech).

Prior to RNA extraction, biomass samples were immediately immersed into RNA*later*^®^ RNA Stabilization Reagent (Thermo Fisher Scientific). Briefly, 50 mL activated sludge was sampled and centrifuged at 3000 rpm for 30 s. The supernatant was pipetted out to leave 1 mL, and the residue was mixed with 10 mL of RNA*later* reagent in 15 mL tubes. The mixture was kept at 4°C for 24 h and then stored at −20°C. Following the manufacturer’s protocol, total RNA was extracted using a FastRNA Pro Blue kit (MP Biomedicals, CA, USA). After depleting rRNA with a NEBNext rRNA Depletion Kit (Bacteria) (New England Biolabs, MA, USA) and confirming DNA concentrations were below the detection limit, fragmented 250-bp RNA was used for preparing complementary DNA sequencing libraries with the MGIEasy RNA Directional Library Prep Set (MGI Tech) following the manufacturer’s protocol. The sequencing was performed with DNBSEQ-G400RS (MGI Tech). DNA and RNA short-read sequencing was conducted by Genome-Lead Co., Ltd (Tokyo, Japan).

### High molecular weight (HMW) DNA extraction and nanopore sequencing

For long-read nanopore sequencing, DNA was extracted from R2 biomass samples in duplicate using a phenol-chloroform method [28]. RNA was removed by RNase, and genomic DNA was precipitated and washed with ice-cold ethanol. Extracted DNA was resuspended in TE buffer and stored at 4°C. The integrity and purity of extracted DNA were checked by 1% agarose gel electrophoresis and a Nanodrop spectrophotometer. The extracted DNA concentration and purity were quantified by a Qubit Assay.

HMW DNA (1 µg) was subjected to library preparation using the ligation sequencing kit SQK-LSK109 (Oxford Nanopore Technologies, Oxford, United Kingdom). Following the manufacturer’s protocol, Nanopore sequencing was performed using a MinION Mk1B instrument (R.9 flow cell, FLO-MIN106; Oxford Nanopore Technologies). Base calling was conducted after sequencing using Guppy software version 5.0.11 (Oxford Nanopore Technologies).

### Metagenomic assembly

Quality control of paired-end short reads (SR) was performed by Fastp (v 0.20.1) [29] with specific parameters (-q 30 -n 20 -t 1 -T 1 −l 30). After quality control, more than 35 million pairs and 31 million pairs of reads were obtained for R1 and R2, respectively. The quality-filtered SR from R1 and R2 samples were assembled individually using Megahit (v1.2.9) with parameters (--k-min 21 –k-max 141 –k-step 12).

Nanopore sequencing produced 970718 sequences, with the longest sequence of 261486 bp and a mean read length of 5981 bp; 100% reads were above Q7, and more than 85% reads were above Q10. The raw reads were directly subjected to assembly using Flye (v2.9-b1768) with the “-meta” setting and the “-nano-hq” mode. The quality-filtered SR were mapped to the assembled contigs of long reads (LR) using Burrows-Wheeler Aligner (BWA)-MEM (v0.7.17-r1188) [30] with default parameters. The resulting sequence alignment and map (SAM) files were converted to binary alignment and map (BAM) files and subsequently sorted using SAMtools (v1.14) [31]. The sorted BAM files were used for polishing the assembly of LR using Pilon v1.24 [32] with the option “—fix bases.”

### Recovery of metagenome-assembled genomes (MAGs)

MAGs from R1 and R2 were recovered as previously described [33] with some modifications. Briefly, the polished contigs of LR assembly were merged with the assembled contigs of paired-end SR (individually for R1 and R2) using the function Merge_wrapper.py in Quickmerge (v0.3) [34] with parameters (-ml 7500 -c 3 -hco 8). The merged results were then processed individually for each sample. Automatic binning was performed using MetaWRAP (v1.3.2) [35] binning module with three genome binners: Concoct [36], Maxbin2 [37], and Metabat2 [38]. Additional binning was done with Vamb (v3.0.2) [39]. The bins from all binning tools were consolidated using the Bin_refinement module (parameters: -c 50 -x 10) of MetaWRAP (v1.3.2). The subsequent steps included extraction of SR and LR for the individual bin, reassembly of LR for each bin and reassembly of each bin, and bin polishing, as described elsewhere [33]. The final bins, MAGs, were selected on the basis of quality (completeness – 5 × contamination > 50) [40] determined by CheckM (v1.1.3).

MAGs from both R1 and R2 were combined and dereplicated using dRep (v3.2.2) [41] (-p 32 −l 2000 -pa 0.90 -sa 0.99 -comp 50 -con 10 -nc 0.1). The processed MAGs were first divided into primary clusters using Mash at a 90% Mash average nucleotide identity (ANI) (specified by -pa 0.90). Each primary cluster was then used to form secondary clusters at the threshold of 99% ANI (-sa 0.99) with at least 25% overlap between genomes. The best MAG was selected within each cluster on the basis of completeness, redundancy, N50 of contigs, and fragmentation. MAG taxonomic assignment was obtained using the “classify_wf” workflow of GTDB-Tk (v2.4.0) [42] with the Genome Taxonomy Database (GTDB) R220 [43].

### Metatranscriptomics processing

The remaining adapter sequences in metatranscriptomic data were trimmed with trim galore (v0.6.6) (https://www.bioinformatics.babraham.ac.uk/projects/trim_galore/). Quality control of paired-end metatranscriptomic reads was then processed using the trimFilterPE function of FastqPuri (v1.0) [44] with specified parameters (--length 150 -q 20 -m 20 -Q ENDS -z yes). The rRNA reads in the quality-trimmed reads were filtered out using SortMeRNA (v4.3.4) [45] with the SILVA database (16S and 23S for bacteria and archaea; 18S and 28S for eukaryotes) and Rfam database (5S and 5.8S).

### Annotation and functional analysis

All MAGs were annotated using Prokka [46]. Metabolic profiles of MAGs were also generated with the METABOLIC-C.pl program in METABOLIC software [47]. The functional marker genes encoding key enzymes responsible for N-cycle were searched from both Prokka annotation and HMM search results of METABOLIC. The markers’ nucleotide and amino acid sequences were extracted using the “Subseq” function of Seqtk (https://github.com/lh3/seqtk). The results were compared, manually curated, *i.e.,* length, locus position, and presence of other genes in the operons, and confirmed by Blastp amino acid sequences against NCBI nr databases. Genes encoding putative high-affinity terminal oxidases (cytochrome bd ubiquinol oxidases *cydAB* and cbb3-type cytochrome c oxidases *ccoNO*) and low-affinity terminal oxidases (caa3-type cytochrome c oxidases *coxAB*) were obtained from MAGs [48].

The *amoA* genes encoding ammonia monooxygenase were not found in any recovered MAGs. To conduct a more comprehensive search of *amoA* gene, HMM profiles of *amoA*-ammonia-oxidizing archaea (AOA) (archaeal *amoA*), *amoA*-ammonia-oxidizing bacteria (AOB) (bacterial *amoA*), and *amoA*-complete ammonia-oxidizing bacteria (comammox) (*Nitrospira* comammox *amoA*) were downloaded from Fungene [49]. Hmmsearch using these profiles was conducted against all contigs (> 500 bp), and the results were checked by Blastp.

Universal single copy marker (SCM) genes were extracted using the fetchMGs tool (available at https://motu-tool.org/fetchMG.html). A subset of 10 SCMs (COG0012, COG0016, COG0018, COG0172, COG0215, COG0495, COG0525, COG0533, COG0541, COG0552) were selected following previous studies [17, 50]. The median transcript abundance of these 10 SCMs was applied to estimate the transcription activity of the MAG [50, 51].

### Calculating relative abundance and mRNA expression

The quality-filtered metagenomic SR and non-rRNA metatranscriptomic reads of each sample were mapped to all dereplicated MAGs using BWA-MEM (v0.7.17-r1188) [30] with the default setting. The unmapped reads were filtered out with SAMtools view (-hbS -F4), and the BAM file was subsequently sorted with SAMtools sort. The coverage of each MAG was calculated using the “genome” mode of coverM (v0.6.1) (https://github.com/wwood/CoverM) with the transcripts per million (TPM) method [52].

To calculate the abundance and mRNA expression of functional marker genes, the reads were mapped against the extracted nucleotide sequences using bowtie2 [53] in “-very-sensitive” mode (to ensure unique mapping). The mapped reads were filtered and sorted as described above. The coverages of marker genes in each sample were obtained by the “contig” mode of coverM (v0.6.1) (https://github.com/wwood/CoverM). The TPM values were normalized to total reads mapped to a sample. The raw-read count was obtained from coverM (v0.6.1) with the “count” method for statistical analysis of the differential mRNA expression using DESeq2 [54].

### Phylogenomic analyses of MAGs and phylogenetic analyses of functional genes

To construct the phylogenetic tree of MAGs, the multiple sequence alignment of 120 bacterial marker genes (*gtdbtk.bac120.user_msa.fasta*) identified and generated by the “classify_wf” workflow of GTDB-tk (v2.4.0) was used to infer the maximum likelihood phylogenetic tree using IQ-TREE2 (v2.2.0) (-T AUTO -m LG). A Newick tree output file was visualized with iTOL v6 (https://itol.embl.de/).

For functional marker genes, amino acid sequences were clustered at a 95% similarity threshold using CD-hit. The corresponding references for each marker are presented in **Table S1**. The amino acid sequence clusters and reference sequences were combined and aligned with Muscle [55] (for the individual key functional genes). The phylogenetic tree was \ with IQ-TREE2 [56] (v2.2.0) (-m MFP -B 1000 -T AUTO). The best model for a phylogenetic tree was selected in accordance with Bayesian information criterion scores and weights (BIC). The tree was visualized with iTOL v6 and used to phylogenetically classify the sequences in this study (**Figs. S2** and **S3**).

### Data depositions

The raw 16S rRNA gene amplicon, metagenomic, and metatranscriptomic sequencing data are available in the DNA Data Bank of Japan (DDBJ) nucleotide sequence database under bioproject accession numbers of PRJDB17920, 17921, and 17922, respectively. Assembled and annotated MAGs have been deposited in the DDBJ nucleotide sequence database with the accession numbers SAMD00776737-SAMD00776788. Metatranscriptomic sequencing data have been deposited with the accession numbers SAMD00770009-SAMD00770012.

## RESULTS

### The dynamics of nitrogen species

The MAS systems were stably operated on day 268, just before the aeration rate was doubled. DO was below the detection limit at an aeration rate of 2 L/min. Gaseous N_2_O in the headspace of R2 was 2.6 ppmv, while NO_3_ concentration ranged from undetectable levels to 0.7 mg-N/L (**Fig. S1**). The increase in the aeration rate to 4 L/min for 15 h surged DO concentration to 6.5 mg/L, in conjunction with a linear increase in NO_3_ concentration of 2.8 mg-N/L (**Fig. S1**). The off-gas N_2_O concentration increased to 6.5–10.3 ppmv, in line with our previous observation [5]. Decreasing aeration to the original level (2 L/min) plunged DO and NO_3_ concentrations. This stepwise change in aeration activated ammonia oxidation at a high DO concentration and increased N_2_O emission when transitioning from anoxic to oxic conditions.

### Microbial community structure

Ninety-seven and one non-redundant MAGs belong to Bacteria and Archaea (genus *Methanomassiliicoccus*), respectively. Among these, 53 MAGs were above the criteria, with completeness > 90% and contamination < 5%; the average completeness and contamination were 85.7% and 1.6%, respectively (**Supplementary Text S1** and **Table S2**). In total, 82.4% and 84.3% of high-quality SR from metagenomic data of R1 and R2 samples, respectively, were mapped to the 98 non-redundant MAGs. Additionally, 84.5 ± 0.01% (n=3) and 82.8 ± 0.00% (n=3) non-rRNA metatranscriptomic reads mapped to 98 MAGs for control and high DO samples, respectively, indicating that the recovered MAGs well represented the microbial community in MAS systems.

In an overall comparison, the microbial profiles by the MAGs and by amplicon sequencing of the V4 16S rRNA region displayed a similar composition of the dominant phyla/class (R1_16S *vs*. R1_MG and R2_16S *vs*. R2_MG in **Fig. 1**). A similar trend of the relative abundance at a phylum level (class for Proteobacteria) between R1 and R2 was captured by both methods (R1_16S *vs*. R2_16S and R1_MG *vs*. R2_MG). Higher relative abundances of Gamma- and Alphaproteobacteria were detected in R2 than in R1, while those of Chloroflexota and Bacillota were higher in R1 than in R2 (**Fig. 1**).

**Figure 1:**
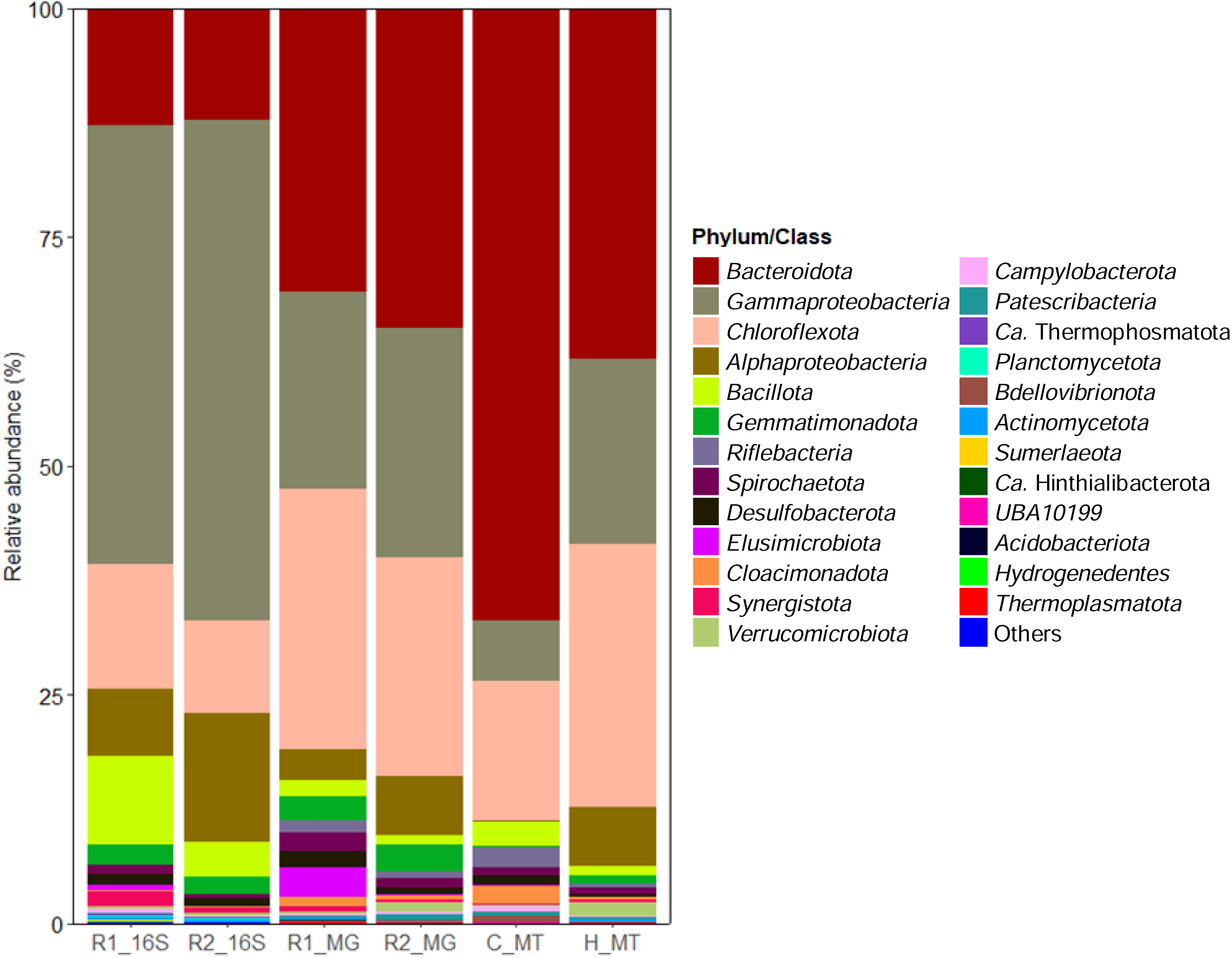
Relative abundance and mRNA expression profiles of microbial community at phylum (class level for Proteobacteria) level in microaerophilic activated sludge systems based on amplicon sequencing of the V4 region in the 16S rRNA gene (R1_16S and R2_16S), metagenome-assembled genomes (MAGs) (R1_MG and R2_MG), and metatranscriptomic reads of reactor 2 (R2) under microaerophilic (C_MT) and high DO (H_MT) conditions mapped to MAGs. Bacterial phyla with abundance < 0.03% were grouped into Others.

In contrast, the 16S rRNA gene data showed higher fractions in Proteobacteria (55.3%–68.7%) and Bacillota (3.8%–9.6%) than MAGs data (24.9%–31.2% for Proteobacteria and 1.1%–1.8% for Bacillota), while the opposite trend was obtained regarding the fractions of Bacteroidota [30.9 %–34.9% (MAG) *vs.* 12.3%–12.7% (16S)] and Chloroflexota [23.8%–28.6% (MAG) *vs.* 10.2%–13.7% (16S)]. Additionally, the metagenomics approach retrieved MAGs affiliated with the families of Ozemobacteraceae, *Candidatus* Hinthialibacteraceae, and JAAZCA01 in three *Candidatus* phyla, Riflebacteria, Hinthialibacterota, and UBA10199, that were not captured by 16S rRNA gene amplicons. The differences are presented in **Text S1**.

### The overall response of microbial activity to DO concentration

The metatranscriptomic data showed that Bacteroidota, which are facultative anaerobes [57], was the most active population in the MAS system, accounting for 66.9% of gene expression under the microaerophilic condition; it decreased to 38.3% under high DO conditions (**Fig. 1**). The second and third transcriptionally active groups were Chloroflexota and Gammaproteobacteria, respectively. The transcription activities were two and three times lower in the microaerophilic condition than in the high DO condition for Chloroflexota [15.2% (control) *vs.* 28.7% (high O_2_)] and Gammaproteobacteria [6.5% (control) vs. 20.3% (high O_2_)], respectively.

The most downregulated population under a high DO condition was Bdellovibrionota (15-fold), followed by Cloacimonadota (5.7-fold). Other bacterial taxa (Riflebacteria, Campylobacterota, and Desulfobacterota) were downregulated by factors of 4.4–4.8. In contrast, increasing DO concentration activated Alphaproteobacteria (26-fold), followed by Actinobacteriota (23-fold), Verrucomicroiota (16-fold), and Gemmatimonadota (9.2-fold).

### The predominant and active bacterial members of the MAS systems

Analyzing the relative abundance and RNA expression of the recovered MAGs allowed for predicting the *in situ* contribution of the community members to the system performance. Overall, the predominant and transcriptionally active bacterial species in the MAS system were identical. An unclassified species (UBA8950 R1_bin.54_o, Chloroflexota) was the most prominent MAG (average log_2_TPM of 17.5) and was the third and the second most active member under control (log_2_TPM of 16.6) and high DO conditions (log_2_FC of 0.8), respectively (**Fig. 2**). The following top abundant MAGs are affiliated with Bacteroidota (in decreasing order of average abundance): unclassified species of UBA6192 (R2_bin.12_r, log_2_TPM of 16.1), *Cloacibacterium* sp. 002422665 (R1_bin.104_o, log_2_TPM of 16.0), and unclassified species of genus *Paludibacter* (R2_bin.117_r, log_2_TPM of 15.7). While the activities of these three Bacteroidota members slightly decreased with increasing DO concentrations, *Cloacibacterium* sp. 002422665 was the most active member in the community under both controls (log_2_TPM of 18.4) and high DO (log_2_TPM of 17.8) conditions. UBA6192 MAG and *Paludibacter* MAG were the top active bacterial species under both conditions (**Fig. 2**). Other abundant bacterial species belong to Gammaproteobacteria (*Rubrivivax, Thermomonas*, and Burkholderiaceae) with an average log_2_TPM of 15.0–15.4. *Thermomonas* MAG was the top active member under both conditions, whereas the transcription activities of *Rubrivivax* (R1_bin.6_o) and Burkholderiaceae (R1_bin.59_o) were moderate under microaerophilic conditions (log_2_TPM of 12.1 and 11.1, respectively) and significantly upregulated (log_2_FC of 2.2 and 3.2, respectively) at a high DO concentration (adj *p-value* < 0.05). The downregulated and upregulated bacterial members at the high DO concentration are provided in **Supplementary Text S2**.

**Figure 2:**
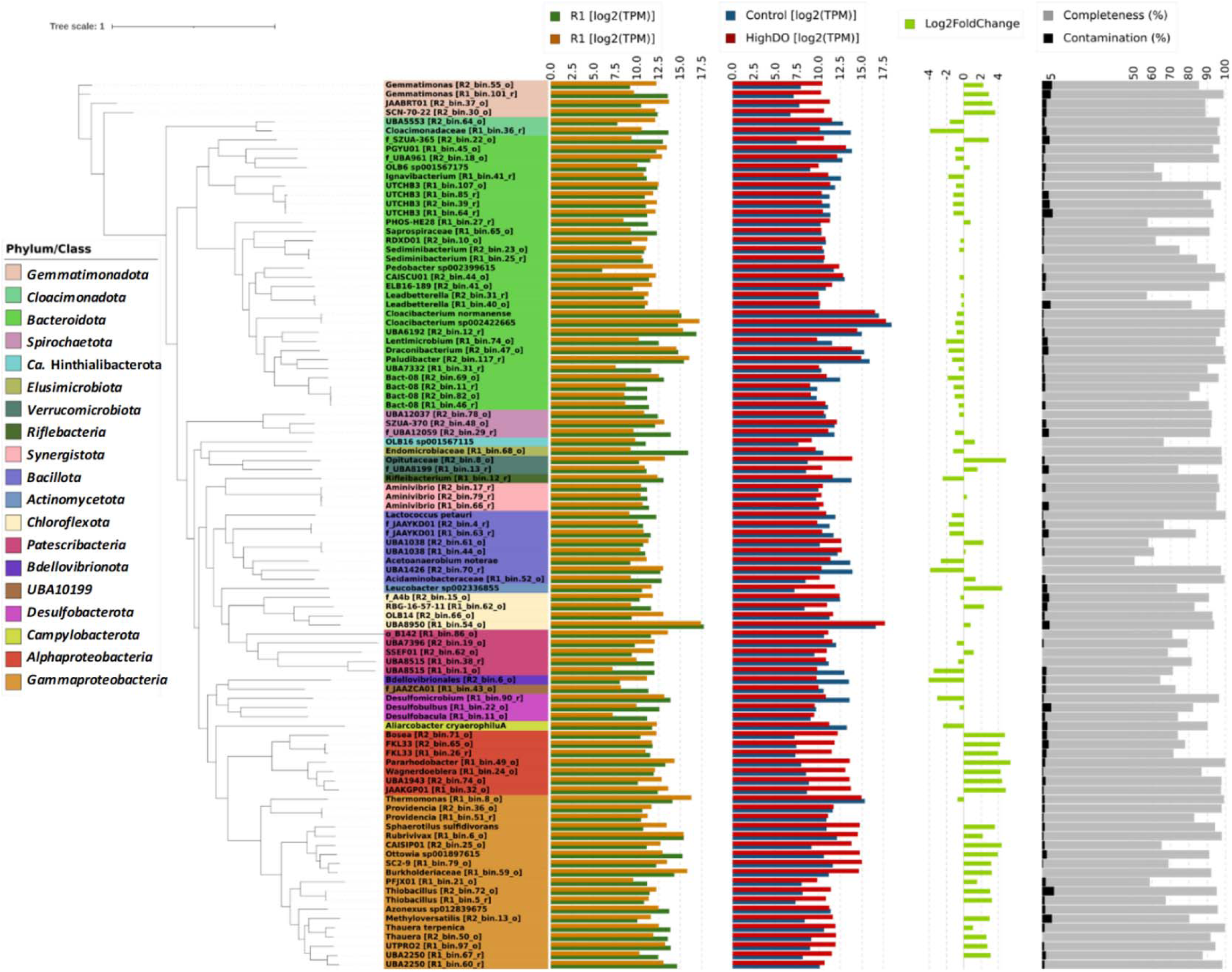
The phylogenomic tree of 97 bacterial metagenome-assembled genomes (MAGs) recovered from this study. The phyla in the tree are colored, whereas the phylum Proteobacteria is further broken down into a class level. The tip labels are the lowest taxonomic classification of MAGs by GTDB-tk (v1.7.0) with GTDB R202. The MAG IDs are included in the square brackets, except for MAGs, which are assigned with species names. The innermost bar plots show the relative abundance (log_2_TPM) of MAGs recovered from metagenomics data of R1 (green) and R2 (orange). The second innermost bar plots represent the average mRNA expression (log_2_TPM) of MAGs in metatranscriptomics data of triplicate biomass samples from R2 under microaerophilic (control, dark blue) and high dissolved oxygen (red) conditions. The next column in light green is the normalized mRNA expression [log_2_ foldchange (High DO/Control)] over the control value as the denominator, calculated by DEseq2. Only statistically different values (adj *p-value* < 0.05) are shown. The outermost bar plots show the contamination (black, maximum value = 6.26%) and completeness (light grey) of MAGs by CheckM. The tree was constructed by IQ-TREE2 (v2.2.0) using multiple sequence alignment of 120 bacterial marker genes generated by “classify_wf” of GTDB-tk.

### Nitrogen metabolism

#### Ammonia oxidation

The MAS system aims to retain ammonia by suppressing ammonia oxidation [5]. As expected, an HMM search using hmm profiles of archaeal, bacterial, and comammox AmoA did not return a hit from the 98 recovered MAGs. A comprehensive search against all assembled contigs (> 500 bp) obtained three hits for both bacterial and comammox AmoA profiles (with each contig < 1000 bp), while no hit was returned for the archaeal AmoA profile. The phylogenomic analysis identified them as *Nitrosomonas* AmoA (**Fig. 3**). The presence of *amoA* genes in the metagenomic data was sporadic, with total abundances of 0.16 TPM (R1) and 0.34 TPM (R2). The full expression under the microaerophilic condition was 0.05–1.01 TPM (**Table 1**). Notably, increasing DO concentration dramatically induced the expression of *amoA* genes with a total expression of 22.7–29.1 TPM, which was an average 57.7 times higher than expression in the microaerophilic condition (**Table 1**). This result highlighted the suppressive effect of the microaerophilic condition on AOMs in the MAS systems. A low amount of AOB (*Nitrosomonas*) was present in the system, while AOA and comammox bacteria were absent.

**Figure 3:**
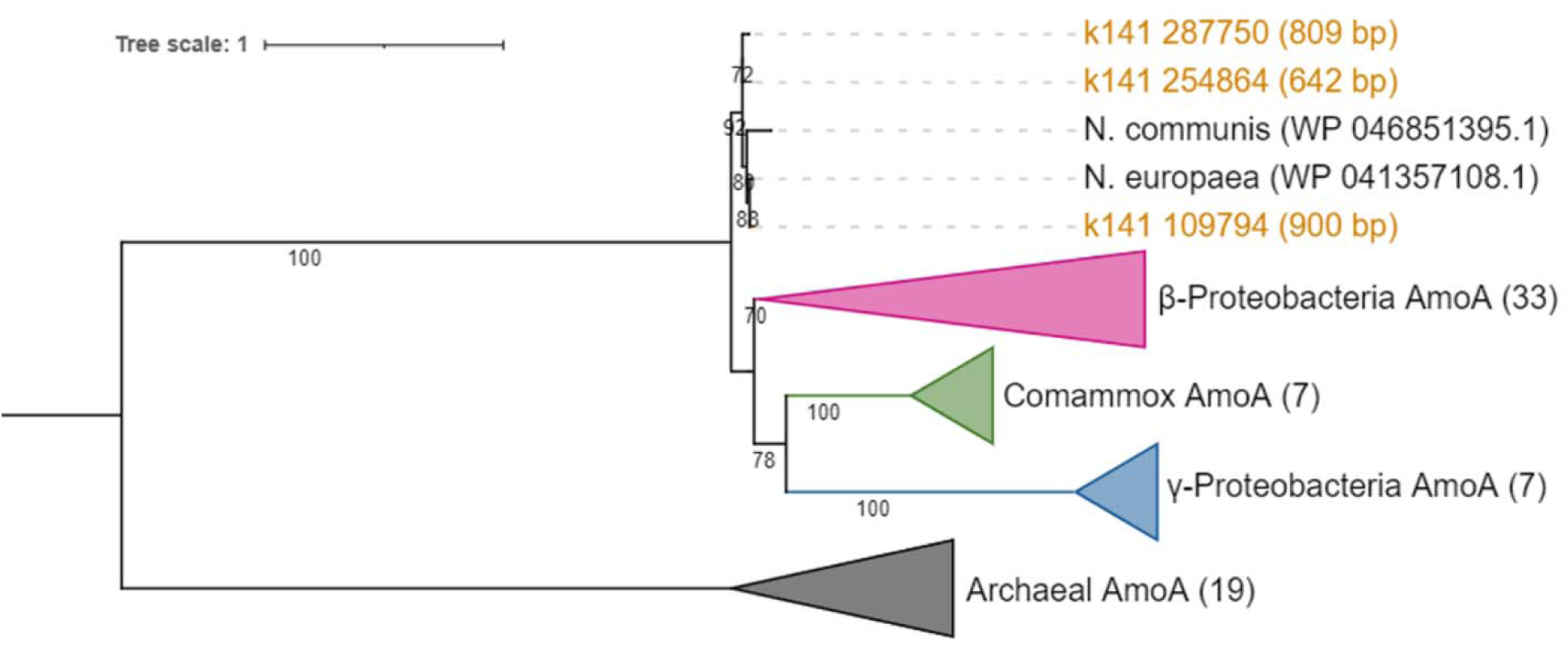
Phylogenetic tree of ammonia monooxygenase subunit A (AmoA) proteins. Amino acid sequences obtained in this study are shown in orange. The lengths of nucleotide sequences (bp) are included in parentheses. The reference AmoA sequences for AOB, Comammox, and AOA were downloaded (seed sequences) from FunGene (http://fungene.cme.msu.edu/, accessed date: 23 March 2022). The number of reference sequences in the collapse clades is indicated in each parenthesis. The bootstrap numbers (1000 replicates) are shown along the branches. The scale bar indicates estimated substitutions of amino acids. The tree was rooted at the midpoint.

**Table 1:** The relative abundance and expression of the *amoA* gene, expressed as TPM (normalized to the gene length and total reads mapped to the sample).

| Contig | Metagenomics |  | Metatranscriptomics |  |  |  |  |  |
| --- | --- | --- | --- | --- | --- | --- | --- | --- |
|  | R1 | R2 | Control<br>1 | Control<br>2 | Control<br>3 | High DO<br>1 | High DO<br>2 | High DO<br>3 |
| k141_254864<br>(642 bp) | 0.03 | 0.00 | 0.04 | 0.07 | 0.00 | 0.11 | 0.15 | 0.22 |
| k141_109794<br>(900 bp) | 0.09 | 0.19 | 0.02 | 0.84 | 0.22 | 17.53 | 20.61 | 19.46 |
| k141_287750<br>(809 bp) | 0.04 | 0.15 | 0.00 | 0.10 | 0.05 | 7.49 | 8.32 | 2.97 |
| Sum | 0.16 | 0.34 | 0.05 | 1.01 | 0.27 | 25.13 | 29.09 | 22.65 |

### Microbial activity of nitrate respiration in the MAS community

All genes involved in nitrogen oxide respiration were found in the recovered MAGs (**Fig. 4A**). The most abundant gene was *narG* (386-405 TPM), followed by *nrfA* (285-366 TPM), *nirS* (241-242 TPM), and clade II *nosZ* (129-254 TPM). The dominant gene types in reduction of nitrogen oxides were *narG* among nitrate reductase (∼5.1 times of *napA*), *nirS* among nitrite reductase (∼3.2 times of *nirK*), *cnorB* among nitric oxide (NO) reductase (∼1.6 times of *qnorB*), clade II *nosZ* among N_2_O reductase (∼9.4 times of clade I *nosZ*), and *nrfA* among DNRA (∼2.9 times of *octR* and ∼19 times of *nirB*). Clade I *nosZ* was the least abundant (18–22 TPM) of all genes for nitrogen oxide reduction.

**Figure 4:**
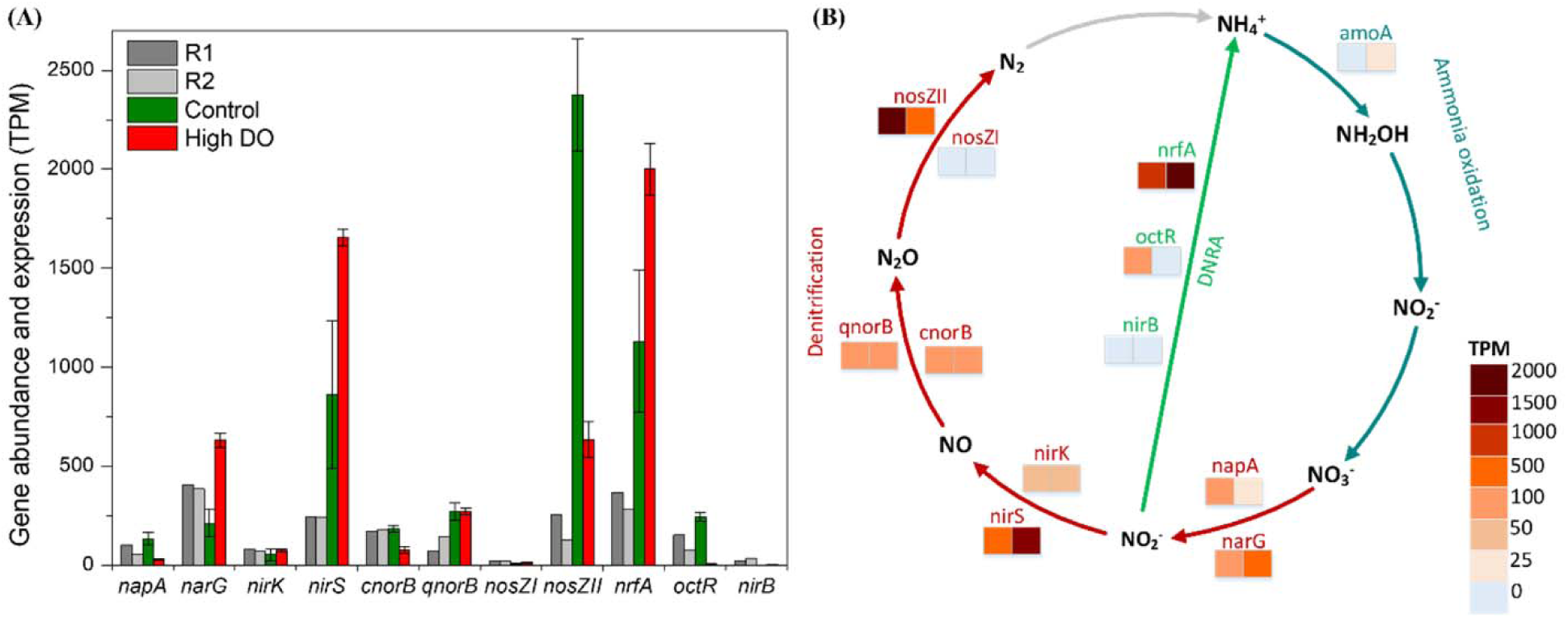
(**A**) Relative abundance (metagenomics R1 and R2) and expression (metatranscriptomics control and high DO) of nitrate respiration genes. Values represent transcripts per million (TPM). Error bars represent the standard deviation of triplicate metatranscriptomic samples. **(B)** A diagram of nitrogen metabolisms in the MAS system: nitrification (dark green), denitrification (dark red), and DNRA (light green). Heatmap color bars indicate the expression level of genes (transcriptomics) with control in the left squares and high DO in the right squares.

As for transcripts, clade II *nosZ* was the most highly expressed gene in the control (microaerophilic) condition (2375 ± 282 TPM, n = 3) (**Fig. 4A** and **B**). The second and third most expressed genes were *nrfA* (1130 ± 359 TPM, n = 3) and *nirS* (861 ± 372 TPM, n = 3), respectively, which were approximately 3.5 times as high as the gene abundances (**Fig. 4A** and **B**). Other gene expressions were low, ranging from 0 (*nirB*) to 269 TPM (*qnorB*). The *narG* transcript was half of its metagenomic abundance under microaerophilic conditions. Clade I *nosZ* and *nirB* genes were scarcely expressed.

Increasing DO concentration significantly repressed the expression of clade II *nosZ* and *cnorB,* which decreased by ∼3.8 times (633 ± 91 TPM, n = 3) and ∼2.5 times (76 ± 17 TPM) in high DO conditions, respectively. *qnorB* gene expression was kept constant (269 ± 44 TPM and 270 ± 17 TPM under microaerophilic and high DO conditions). In stark contrast, *narG*, *nirS*, and *nifA* gene expressions increased by ∼3.0, 1.9, and 1.7 times under high DO conditions, respectively (**Fig. 4A** and **B**). While clade I *nosZ* expression doubled at high DO concentrations, the total expression was negligible (11 ± 4 TPM, n = 3) (**Fig. 4A**).

### Microbial sinks of N_2_O in MAS systems

Of the 98 recovered MAGs, 73 MAGs (>74%) from 11 bacterial phyla harbor genes involved in nitrate respiration. Of the 73 MAGs, 39 MAGs (∼53%) carried *nosZ* genes. The phylogenetic allocation at a genus level is shown in **Fig. S2**. Notably, 31 MAGs harbor clade II *nosZ* and were distributed mainly in Bacteroidota (22 MAGs), followed by Gammaproteobacteria (5 MAGs), Gemmatinomadota (3 MAGs), and Chloroflexota (1 MAG) (**Fig. 5**). Only 8 MAGs carried clade I *nosZ* and consisted exclusively of Alpha-(4 MAGs) and Gammaproteobacteria (4 MAGs). All Alphaproteobacteria MAGs carrying clade I *nosZ* possessed only *nirK* and *cnorB* as denitrifying genes. These MAGs expressed low levels of *nirK* (3.1–5.3 TPM) and generally did not express other genes (**Fig. 5**).

**Figure 5:**
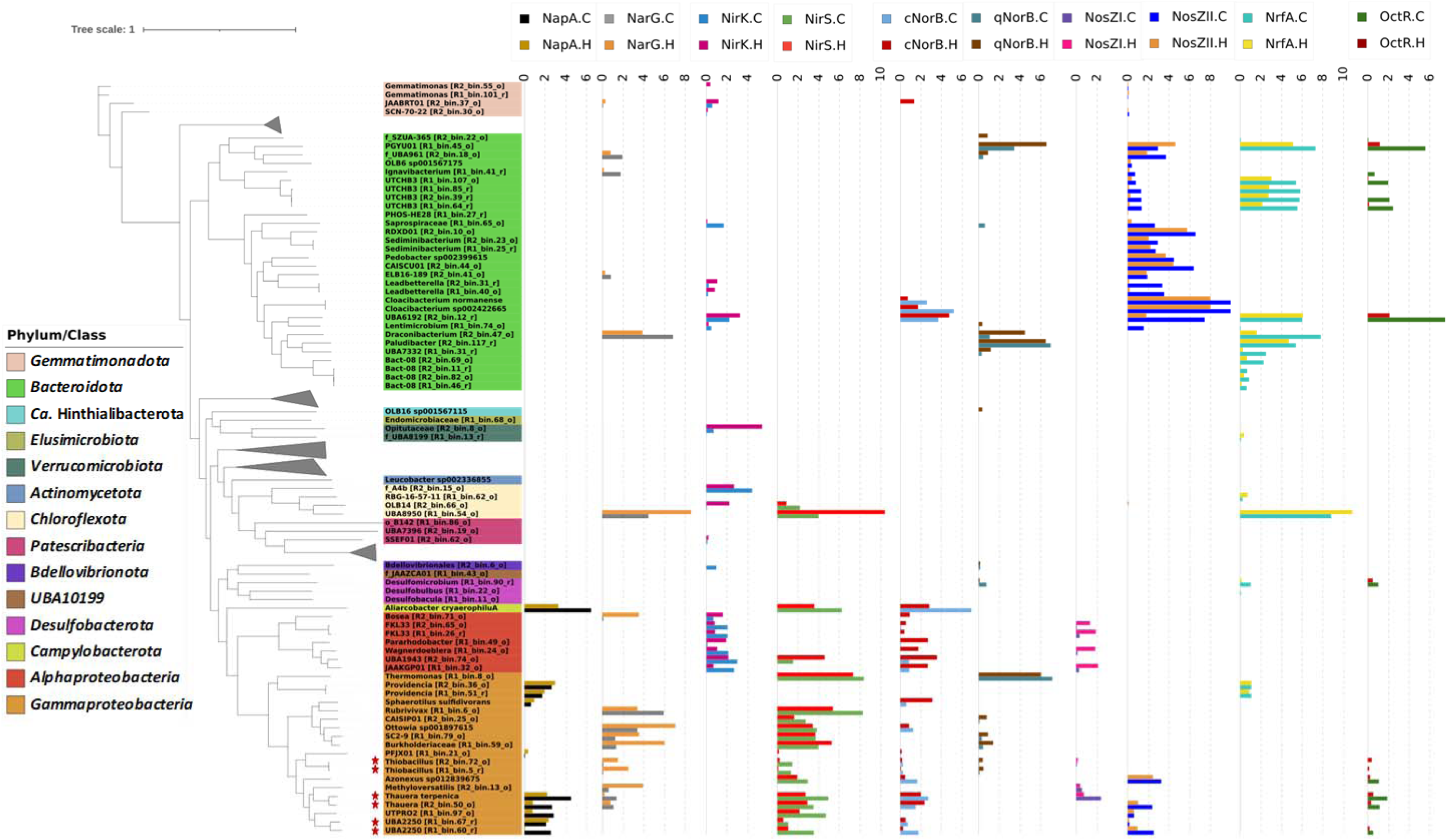
The expression of nitrogen cycle-associated genes of metagenome-assembled genomes (MAGs) obtained from a MAS system. The gene expression was expressed as log_2_(TPM+1). The “*” indicates MAGs possessing all denitrifying genes (canonical complete denitrifying bacteria).

Six MAGs possessed a complete set of denitrification genes, all belonging to the order Burkholderiales (Gammaproteobacteria). These canonical complete denitrifiers harbor *nirS* type (no *nirK* type) and *cnorB* (except for *Thiobacillus* harboring both *cnorB* and *qnorB*). Three complete denitrifying MAGs, *i.e.,* two unclassified *Thiobacillus* species (R1_bin.5_r and R2_bin.72_o) and *Thauera terpenica* (R2_bin.16_o), carried clade I *nosZ*. The other three complete denitrifying MAGs carried clade II *nosZ* (two unclassified UBA2250 species [R1_bin.60_r and R1_bin.67_r] and one unclassified *Thauera* species [R2_bin.50_o]). Notably, the six complete denitrifying MAGs do not have *nrfA* genes, but all contain *octR* genes for DNRA. Regardless of the clade type, all denitrifying genes in these MAGs were expressed at low levels.

Among 31 MAGs harboring clade II *nosZ*, 18 MAGs were non-denitrifying bacteria missing genes encoding nitrate and nitrite reductases (**Fig. 5**). Although the expression level decreased from a microaerophilic to a high DO condition (970 TPM to 250 TPM), the highest expressions levels of clade II *nosZ* were found in two *Cloacibacterium* species (*C. normanense* [R1_bin.29_r] and *Cloacibacterium* sp. 002422665 [R1_bin.104_o]) in both conditions. These two species harbor the *cnorB* gene but no nitrate and nitrite reductase genes (**Fig. S4**). The *cnorB* gene expressions of *Cloacibacterium* sp. 002422665 and *C. normanense* were suppressed at a high DO concentration (< 2 TPM).

The other two non-denitrifying MAGs with the top expressions of clade II *nosZ* were RDXD01 MAG [R2_bin.10_o] and CAISCU01 MAG [R2_bin.44_o] in Bacteroidota (**Fig. 5**). These MAGs possess only *nosZ* gene among denitrifying genes. *nosZ* expression decreased from 92.3 ± 20.9 to 51.8 ± 6.8 TPM (RDXD01 species) and 80.9 ± 9.4 to 20.1 ± 4.0 TPM (CAISCU01 species) upon switching from microaerophilic to high DO conditions.

The uncharacterized UBA6192 species (R2_bin.12_r) in Bacterioidota, possessing clade II *nosZ*, *nirK*, *cnorB*, and DNRA genes (*nrfA* and *octR*), showed the unique gene expression patterns. It displayed marginal expressions of *nirK* and *cnorB* genes. Introducing a high DO condition from the microaerophilic condition suppressed the gene expressions of *nosZ* and *octR*, whereas maintaining the gene expression level of *nrfA* and increasing *nirK* and *cnorB* gene expressions (8.5 ± 0.3 TPM and 25.0 ± 2.5 TPM, respectively).

### Microbial source of NO and N_2_O in MAS systems

*Thermomonas* (R1_bin.8_o, Gammaproteobacteria), *Paludibacter* (R2_bin.117_r, Bacteroidota), and *Aliarcobacter* (R2_bin.54_o, Campylobacterota) devoid of *nosZ* genes displayed high expressions of *norB* genes, likely ascribed to N_2_O sources in the MAS system. *Thermomonas* showed the highest expressions of both *nirS* (328.4 ± 99.1 TPM) and *qnorB* (134.8 ± 36.1 TPM) under microaerophilic conditions. Both gene expressions decreased by half under high DO conditions (157.3 ± 11.5 TPM for *nirS* and 62.8 ± 5.2 TPM for *qnorB*). *Paludibacter* species possessed only the *qnorB* gene among denitrifying genes, with high expression (121.1 ± 5.3 TPM), and *nrfA* gene, with moderate expression (40.1 ± 4.4 TPM), under microaerophilic conditions. Both *qnorB* and *nrfA* genes were slightly suppressed (∼1.5 times) under high DO conditions.

The potential contribution to the microbial NO sources in the MAS system stemmed from the activities of *Rubrivivax* (R1_bin.6_o, Gammaproteobacteria) and UBA8950 (R1_bin.54_o, Chloroflexota). Both species carried only *narG* and *nirS* among the denitrifying genes. Under the microaerophilic condition, *Rubrivivax* species showed high expressions of *nirS* (309.2 ± 181.2 TPM) and *narG* (58.5 ± 42.5 TPM); these levels were noticeably suppressed by 87% and 68%, respectively, after switching to a high DO condition. In contrast, *nirS* and *narG* gene expressions of UBA8950 species were low under microaerophilic conditions (14.7 ± 13.6 TPM and 20.4 ± 15.1 TPM, respectively) and significantly increased (1344.4 ± 26.1 TPM and 371.7 ± 22.8 TPM, respectively) at high DO concentrations. UBA8950 displayed the same transcription trend of *nrfA* gene, which is addressed in the later section.

### DNRA

Among the 73 MAGs carrying nitrate respiration-related genes, 24 MAGs (∼33%) possessed the *nrfA* gene; this group mainly comprised Bacteroidota (15 MAGs); others belong to Desulfobacterota (3 MAGs), Chloroflexota (2 MAGs), Gammaproteobacteria (2 MAGs), Gemmatinonadota (1 MAG), and Verrucomicrobiota (1 MAG). The phylogenetic allocation at a genus level is shown in **Fig. S3**. Notably, all 8 MAGs harboring the clade I *nosZ* did not carry the *nrfA*, while 8 MAGs (∼26%) of 31 MAGs harboring clade II *nosZ* also carried *nrfA*. Among the claded II *nosZ* MAGs, UBA8950 (R1_bin.54_o, Chloroflexota) actively expressed *nrfA* most. As shown in **Figs. 2** and **5**, this MAG was the most abundant species and harbored *narG* and *nirS* genes. Increasing a DO concentration dramatically upregulated *nrfA* expression, from 445.5 ± 335.5 TPM (microaerophilic condition) to 1849.0 ± 172.3 TPM (high DO concentration), together with the upregulation of *nirS* and *narG* genes (**Fig. 5**). This trend was exclusively observed among the MAGs possessing both *nirS* and *nrfA* for nitrite reduction.

The active DNRA members under microaerophilic conditions were Bacteroidota. *Draconibacterium* (R2_bin.47_o) and PGYU01 (R1_bin.45_o) had high *nrfA* expressions of 222.0 ± 9.9 TPM and 154.2 ± 15.9 TPM, respectively. The other six MAGs of Bacteroidota (one UBA6192, four UTCHB3, and one Paludibacter) expressed *nrfA* in the range of 40.1–62.4 TPM (**Fig. 5**). Except for UBA6192, *nrfA* expressions from all these MAGs were downregulated under high DO conditions.

Of 15 MAGs with *octR* genes, only 2 (UBA6192 and PGYU01) significantly expressed *octR* gene (178.7 ± 26.7 TPM and 46.3 ± 10.0 TPM, respectively). *octR* expressions were downregulated under a high DO condition. While six Gammaproteobacteria MAGs with an entire set of denitrifying genes harbored *nirB* gene, they did not express *nirB* under both examined conditions.

### Growth strategy of microorganisms in MAS systems

The transcripts of genes encoding terminal oxidases, clade II nosZ, and nrfA genes (Fig. 6) were compared to characterize the growth strategies of microorganisms in the MAS system. Briefly, the transcription patterns were diverse. *Cloacibacterium* sp. 002422665 (R1_bin.104_o) showed noticeable expression of high-affinity terminal oxidase *ccoNO* (> 12 times of SCM) and clade II *nosZ* (7 times of SCM) under the microaerophilic condition (**Fig. 6**). The increase in the DO concentration downregulated both *ccoNO* and clade II *nosZ* gene expressions by 2.6 and 3.8 times, respectively. Bacteroidota UBA6192 (R2_bin.12_r) showed higher expressions of clade II *nosZ* and *octR* genes than *ccoNO* under microaerophilic yet not aerobic conditions (**Figs. 5** and **6**). Meanwhile, the expression levels of *nrfA* and *ccoNO* remained relatively constant. Chloroflexota UBA8950 (R1_bin.54_o), possessing *cydAB* devoid of *ccoNO*, upregulated high-affinity terminal oxidase *cydAB*, *nrfA*, and *nirS* genes under a high DO condition, with the SCM expression increased by > 7.0 times (**Figs. 5** and **6**). A higher gene expression of low-affinity oxidases (*coxAB*) and *narG* than SCM was also observed after increasing DO concentration (**Figs. 5** and **6**). The expression patterns of other active members are described in **Supplementary Text S3**.

**Figure 6:**
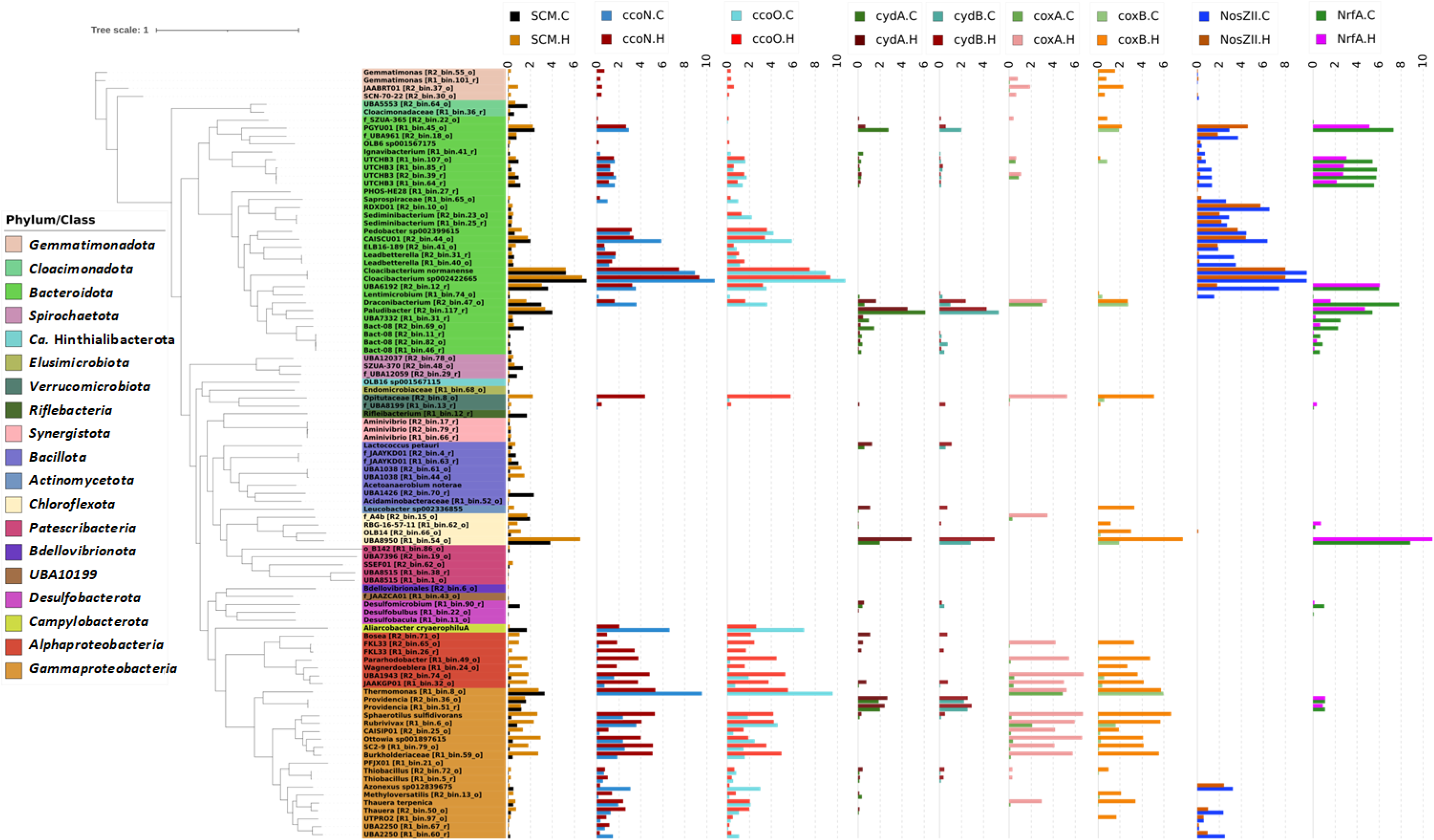
Transcript abundances of high-affinity terminal oxidases (*ccoNO* and *cydAB*) and low-affinity terminal oxidases (*coxAB*) in comparison with transcript abundances of clade II *nosZ* (NosZII), *nrfA*, and single copy markers (SCM). “C” and “H” represent control and high DO conditions, respectively. The gene expression was expressed as log_2_(TPM+1). The TPM value of SCM is the median TPM of the 10 selected SCMs.

## DISCUSSION

A profound understanding of the microbial community and activity in a MAS system is critical for optimizing the system design and operation for innovative ammonia retention and recovery [5]. This study used hybrid sequencing of high-quality Illumina short-reads and Nanopore long-reads and recovered MAGs to unravel the phylogeny, functions, and transcriptomic activities of MAS communities. Metagenomics and metatranscriptomics analyses supported our hypotheses, in which: (1) N_2_O reduction by non-denitrifying bacteria was compromised under high DO conditions, causing N_2_O emission likely by AOB; (2) clade II *nosZ* non-denitrifying bacteria (mainly Bacteroidota) were responsible for N_2_O reduction under microaerophilic conditions; and (3) 26% of clade II *nosZ* bacteria (8 out of 31 MAGs) harbor *nrfA*, potentially exerting N_2_O consumption and DNRA. We further revealed transcriptionally active bacteria driving nitrogen conversions, their functional roles in nitrogen metabolisms, and transcriptional responses to changes in DO concentration in the MAS system (**Fig. 7**).

**Figure 7:**
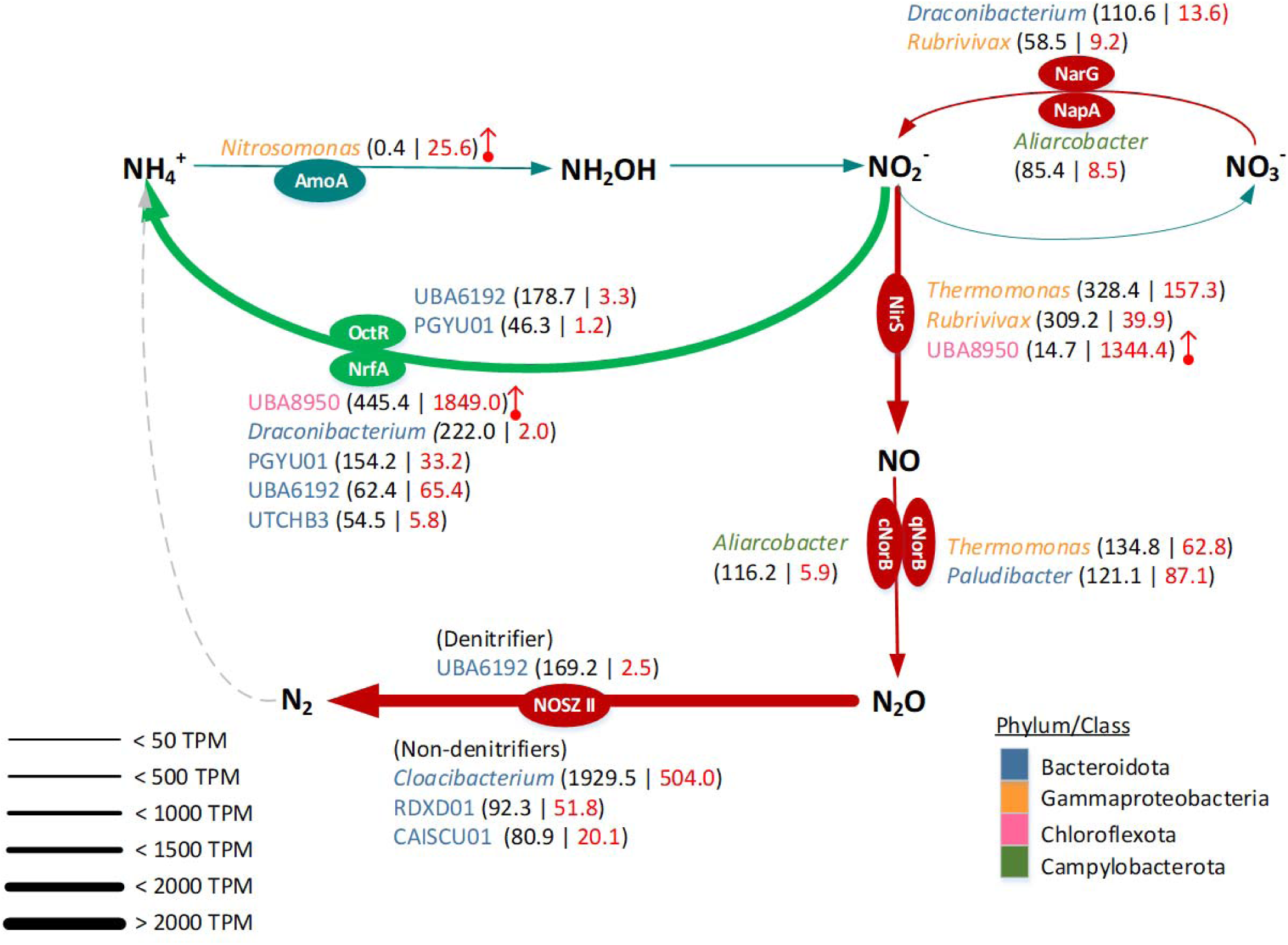
Diagram illustrating the contribution of pivotal microbial members to nitrogen transformations in the MAS system. Three key pathways, nitrification (dark green), denitrification (dark red), and DNRA (light green), with the key enzymes detected in metatranscriptomic data, are shown. The thickness of each continuous arrow indicates the transcription level of an assigned metabolic pathway of nitrogen transformation in the community. The microbial members responsible for each pathway are appended and colored according to phylum/class. The numbers in parentheses show the transcripts (> 45 TPM) of the key enzymes from each microbial member under control (left in black) and high DO (right in red) conditions. The remarkable gene upregulation under a high DO condition is highlighted by a vertical red arrow.

### Identification of AOM and amoA gene expression

No AOM MAG was recovered from the community, and metagenomic data confirmed the rare existence of the *Nitrosomonas amoA* gene in the MAS system. Operation under the microaerophilic condition successfully inhibited *amoA* expression; however, increasing DO concentration dramatically upregulated *amoA* expression, resulting in ammonia oxidation, congruent with the increase in nitrate concentration (**Fig. S1**). This finding agrees with AOB detection (but unnoticeable respiratory activities) by 16S rRNA amplicon sequencing, qPCR, and fluorescence *in situ* hybridization [5]. Hence, precise control of DO concentration was the key to retaining ammonia in the MAS system.

### Genotypes and transcriptions of N_2_O-reducing bacteria

The highest expression of clade II *nosZ* gene among nitrogen metabolism genes (**Fig. 4A**) highlighted the vital role in reducing N_2_O under the microaerophilic condition (**Fig. 7**). This observation corroborates low or marginal N_2_O emissions from the MAS system [5]. Crucial roles of clade II *nosZ* bacteria in N_2_O reduction in WWTPs were previously reported [10, 58]. Repressing clade II *nosZ* gene expression at an elevated DO concentration likely explains the increased N_2_O emissions observed in this study (**Fig. S1**) and our previous work [5].

The metatranscriptomics disclosed non-denitrifying bacteria belonging to Bacteroidota, particularly *Cloacibacterium*, as major N_2_O sinks in the MAS system (**Figs. 5** and **7**). The *Cloacibacterium* highly expressed high-affinity terminal oxidase (*ccoNO*) and clade II *nosZ* (**Figs. 5** and **6**). *ccoNO* expression is instrumental in efficiently scavenging oxygen under microaerophilic conditions, which conserves energy for bacterial growth and protects NosZ from oxygen inhibition. The potential *nosZ* gene expression mechanism is congruent with a previous work reporting oxygen uptake by oxygen-tolerant N_2_O reducers [18]. Downregulation of both *ccoNO* and clade II *nosZ* genes in these non-denitrifiers at a high DO concentration indicated compromising N_2_O reduction at an elevated DO concentration (**Fig. 7**). The phylogenetic tree of *Cloacibacterium* species shows two distinctive branches; one contains the MAGs recovered in this study (*Cloacibacterium* sp002422665 and *C. normanense*) and the other comprises *Cloacibacterium* sp002337005 and *C. rupense* (**Fig. S4A**). The branch covering the two MAGs in this study possesses *cnorB* and clade II *nosZ*, whereas the other branch (*Cloacibacterium* sp002337005 and *C. rupense*) misses *norB* (**Fig. S4B**). All *Cloacibacterium* species possess high-affinity terminal oxidase *cbb3*-type, while *C. rupense*, *C. caeni_A* (*a.k.a.* CB-01), *C. caeni_B* (CB-03) [59], and *Cloacibacterium* sp002337005 additionally possess *cydBD*-type (**Fig. S4B**), likely harboring the nature of oxygen tolerance.

*Cloacibacterium* is promising for N_2_O mitigation in WWTPs. In a fed-batch experiment, *Cloacibacterium* was enriched at a low N_2_O concentration (20 ppmv in supplied gas) with a high abundance, followed by *Flavobacterium* (Bacteroidota), while their abundances decreased at higher N_2_O concentrations [10]. The recently isolated *C. caeni_B* [59] showed a low growth rate and affinity for N_2_O but long-lasting persistence in soils despite the disadvantageous biokinetics [15]. Non-denitrifying Bacteroidota were also predominant N_2_O reducers in full-scale Danish WWTPs [58, 60], in which N_2_O concentration was below 1 mg/L. Additionally, uncultured *C. normanense* was enriched in anaerobic pig manure digestate supplied with N_2_O [61]. No nitrate reduction was confirmed for the isolates of *C. normanense* NSR1 [62], *C. rupense* [63], and *C. caeni* strain B6 [64]. Therefore, *Cloacibacterium* species are likely persistent non-denitrifying N_2_O sinks in the MAS system where limited amounts of N_2_O and NO_3_ are available (**Fig. S1**). A follow-up physiological investigation on the N_2_O affinity of *Cloacibacterium* species isolated from the system will clarify this claim.

### DNRA bacteria and interaction with N_2_O consumers

The high *nrfA* gene expression found in this study likely facilitated the conversion of nitrite into ammonia, resulting in high ammonia retention under microaerophilic conditions. We show that *Chloroflexota* UBA8950 MAG (R1_bin54.o) exceptionally upregulated *nrfA* gene after increasing DO concentration (**Fig. 5**), concomitant with the upregulation of *nirS*, *narG*, low-affinity terminal oxidase (*coxB*), and high-affinity terminal oxidase (*cydAB*) genes (**Figs. 5** and **6**). This expression pattern indicated that UBA8950 utilized different electron acceptors adaptable to alternating redox conditions. The genus UBA8950 currently comprises only two species (GTDB database, v207 Feb 2023) with five draft genomes (four for sp001872455 and one for sp002840685). All genomes were derived from groundwater metagenomes without the culture representative [40]. The UBA8950 MAG in this study was classified as a new species within this genus. The phylogenetic tree position of this MAG lies in the same branch with sp002840685 (**Fig. S5A**). Functional annotation showed that all UBA8950 genomes possess high-affinity (*cydBD*) and low-affinity (*cox*) oxidases. The UBA8950 MAG and sp002840685 genomes carry genes in nitrate reduction, nitrite reduction, and DNRA pathways, whereas genomes of sp001872455 do not (**Fig. S5B**). Isolation and physiological characterization of UBA8950 bacteria are required for the practical application of the MAS system.

Another notable finding in this study is the coexistence of N_2_O reduction and DNRA pathways in the recovered MAGs. Approximately 26% of MAGs possessing clade II *nosZ* gene also harbor the *nrfA* gene, congruent with the previous report [8]. The high expression of the clade II *nosZ* gene and DNRA genes (*nrfA* and *octR*) by an unclassified UBA6192 species (R2_bin12.r) in Bacteridota (**Fig. 5**), taxonomically close to sp002421995 (**Fig. S6A**), suggests the contribution to N_2_O reduction and DNRA in the MAS system. The *nrfA* expression remained unchanged at an elevated DO concentration, contributing to DNRA even under aerobic conditions. Despite carrying *nirK*, *cnorB*, and high-affinity *cbb3*-oxidases genes, the low gene expressions implied a minor contribution to N_2_O production, underscoring that the UBA6192 MAG (R2_bin12.r) is an ideal candidate for improving N_2_O mitigation and ammonia recovery, *i.e.*, reducing nitrate to ammonia. There are 17 draft genomes of 11 species in the GTDB database for the uncultured UBA6192 genus. All 11 species have the nitrite reduction pathway, and 7 species harbor DNRA and/or NO reduction, whereas only 2 species (sp018816765 and sp018336125) possess *nosZ* (**Fig. S6B**), showcasing the genotype uniqueness of UBA 6192 MAG (R2_bin12.r).

Another potential candidate to facilitate ammonia recovery is PGYU01 MAG (R2_bin45.o), displaying higher expressions of both *nrfA* and *octR* genes before increasing a DO concentration (**Fig. 5**). The downregulation of *nrfA* and *octR* genes and upregulation of *qnorB* and clade II *nosZ* genes at a high DO concentration suggests the switching function from DNRA to N_2_O reduction. Comparably high *nrfA* and *octR* gene expressions in UTCHB3 (Ignavibacteriaceae) bacteria, four MAGs recovered in this study with the genotypes allowing N_2_O reduction and DNRA (**Fig. S7B**), indicate an active involvement in DNRA. Although these bacteria did not pronouncedly express *nosZ* compared to PGYU01 MAG, all UTCHB3 showed moderate *nrfA* expression (**Fig. 5**), potentially instrumental in reducing nitrite to ammonia. PGYU01 and UTCHB3 may support high ammonia recovery by reducing nitrite to ammonia when excessive aeration unwantedly enhances nitrification in the MAS system.

### N-converting microorganisms in the presence of DO

The MAS system, supplied with high-strength nitrogenous wastewater, is rate-limited by oxygen supply. Increasing oxygen supply enhances organic carbon removal and, as a downside, initiates ammonia oxidation in conjunction with N_2_O production by AOMs, as shown in **Fig. S1**. Exploring and harnessing bacteria exerting N_2_O reduction and DNRA potentially paves the way towards high ammonia retention with low N_2_O emissions. Our metatranscriptomic analysis allowed the classification of several phenotypical responses to an increase in DO concentration (**Fig. 7**).

The predominant bacterial members responsible for N_2_O reduction and DNRA exhibited high gene expressions of high-affinity terminal oxidases (*ccoNO* or *cydAB*) and nitrogen oxide reductases. However, the preferential electron acceptor and the strategy for growth seem diverse. The concomitant decreases in high-affinity terminal oxidases *ccoNO* and clade II *nosZ* gene expressions of *Cloacibacterium* (**Figs. 5** and **6**) suggest that the effectiveness as an N_2_O sink could be limited under high DO conditions. The downregulations of clade II *nosZ* and *octR* and the unchanged regulation of *nrfA* and *ccoNO* at a high DO concentration, observed in Bacteroidota UBA6192, indicate that N_2_O reduction and DNRA are compromised after increasing DO concentrations. Much higher expressions of *nrfA* and *nirS* genes than *coxAB* and *cydAB* genes of Chloroflexota UBA8950 at a high DO concentration allude to utilizing nitrogen oxides as the primary electron acceptors and facilitating DNRA at a high DO concentration. Reportedly, DNRA, dependent on NrfA activity, proceeds under oxygen-limited conditions [65], although it may be less oxygen-sensitive than anticipated [66]. The highly expressed terminal oxidase gene (**Fig. 6**) may scavenge oxygen in the periplasm of Chloroflexota UBA8950. The underlying mechanism and the occurrence of DNRA warrant future investigations.

By combining online monitoring of the MAS system, metagenomic, and transcriptomic analyses, this study identified transcriptomically active bacteria responsible for high ammonia retention efficiency and low N_2_O emissions in transitioning from microaerophilic to aerobic conditions. Our analysis revealed non-denitrifying N_2_O-reducing bacteria affiliated with clade II *nosZ Cloacibacterium* in the phylum Bacterioidota as a promising N_2_O sink in microaerophilic conditions. While *Cloacibacterium* decreased transcriptomic activities by transitioning from a microaerophilic to an aerobic condition, bacteria responsible for DNRA, *e.g.*, the genus UBA8950 (Chloroflexota), became active, likely switching from N_2_O reduction to DNRA as a primary nitrogen transformation function of the microbial community at elevated oxygen levels. Switching such metabolic functions in the community of the MAS system caused by oxygen level transitions allowed high ammonia retention with fewer N_2_O emissions even during dynamic oxygen fluctuations, which occasionally occurs in an engineered system for nitrogen management. Our findings may help achieve stable operation for ammonia recovery, a new paradigm that turns nitrogen removal into nitrogen recovery.

## ACKNOWLEDGEMENTS

This paper was based on results obtained through the project of JPNP18016 commissioned by the New Energy and Industrial Technology Development Organization (NEDO). We thank Gabrielle White Wolf, PhD, from Edanz (https://jp.edanz.com/ac) for editing a draft of this manuscript.

## DATA AVAILABILITY STATEMENT

The data of this study will be available upon request.

