## Supplementary information for "Meta-omic insights into active bacteria mediating N_2_O mitigation and dissimilatory nitrate reduction to ammonium in an ammonia recovery bioreactor"

**Text S1: Merits of metagenomics and metatranscriptomics approaches**

A hybrid sequencing approach with high-quality Illumina SR and Nanopore LR recovered 98 non-redundant MAGs (dereplication at 99% ANI), meeting the GTDB database criteria described in **2.5**. Only one recovered MAG belongs to Archaea (genus *Methanomassiliicoccus*); 97 non-redundant MAGs belong to Bacteria. Of them, 53 MAGs were above the criteria, having completeness > 90% and contamination < 5%. Average completeness and contamination were 85.7% and 1.6%, respectively. Fourteen MAGs were recovered with fewer than 10 contigs, and three of these MAGs contained only one contig (**Table S2**).

The 16S rRNA gene amplicon is a common method for profiling bacterial communities. However, this method often overestimates some microbial groups and *vice versa* for others, which is noted in this study and previous works (Xu et al., 2021). For example, a higher fraction of Firmicutes and a lower fraction of Chloroflexota in the 16S rRNA profiles than in metagenomic profiles were consistently observed. A low copy number of the 16S rRNA gene in Chloroflexota (1.4 ± 0.7) and a high copy number of the 16S rRNA gene in Firmicutes (6.9 ± 2.9) (https://rrndb.umms.med.umich.edu/) possibly explain for this difference. On the other hand, consistent with this study, a high number of MAGs belonging to Bacteroidota were reconstructed from activated sludge (Singleton et al., 2021), while the 16S amplicon study often observed a much higher abundance of Proteobacteria at a global scale (Wu et al., 2019). Moreover, bacterial groups lack culture representative (named "microbial dark matter") and were at low presence in the reference database, thus not well captured by the 16S amplicon approach. For example, members of Riflebacteria, *Candidatus* Hinthialibacterota , and UBA10199 were only obtained in MAG profiles, but not in 16S amplicon profiles. Finally, the discrepancies between the functional potential in metagenomic data and the actual expression in metatranscriptomic data in this study indicate the importance of combining both metagenomic and metatranscriptomic for studying the functions of the microbial community under specific conditions.

**Text S2: The predominant and active bacterial members of the MAS system**

Consistent with the community-wide observation, the most downregulated bacterial members under the high DO condition were unclassified MAGs of Bdellovibrionales (R2_bin.6_o) and

Cloacimonadaceae (R1_bin.36_r). The abundances of these MAGs were low ($log_2$TPM of 9.5 for Bdellovibrionales MAG) and moderately low ($log_2$TPM of 12.1 for Cloacimonadaceae MAG). Their transcriptomic activities were moderately high under the microaerophilic condition ($log_2$TPM of 13.5 [Bdellovibrionales] and 13.7 [Cloacimonadaceae]) and significantly decreased ($log_2$FC of -4.1 [Bdellovibrionales] and $log_2$FC of -3.9 [Cloacimonadaceae]) at a high DO condition. Unclassified MAGs of UBA1426 (Firmicutes), UBA8515 (Patescibacteria), and *Desulfomicrobium* (Desulfobacterota) displayed the comparable transcriptomic trend (**Fig. 2**).

The most upregulated member at the high DO concentration was an unknown species of *Pararhodobacter* (R1_bin.49_o) in Alphaproteobacteria. This member was moderately abundant ($log_2$TPM of 13.8) but showed low activity under the microaerophilic condition ($log_2$TPM of 7.9). Its relative activity significantly increased ($log_2$FC of 5.4) under the high DO condition. The 10 most upregulated members ($log_2$FC of 3.9 to 4.9) under the high DO condition include one MAG in Verrumicrobiota (Opitutaceae), six MAGs in Alphaproteobacteria (JaaKGP01*, Bosea,* UBA1943*, Wagnerdoeblera,* FKL33 genera), one MAG in Actinomycetota (*Leucobacter* sp. 002336855), and two MAGs in Gammaproteobacteria (CAISIP01 and *Ottowia* sp001897615) (**Fig. 2**).

**Test S3: Additional microorganisms harboring genes of terminal oxidases**

Except for *Cloacibacterium* sp. 002422665 (R1_bin.104_o), Bacteroidota UBA6192 (R2_bin.12_r), and Chloroflexota UBA8950 (R1_bin.54_o), several bacterial members actively expressing terminal oxidase and nitrogen metabolism genes were detected. *C. normanense* (R1_bin.29_r) was the second active member with lower gene expression (mean SCM of 3.7 times lower and *ccoNO* of 3.5 times lower than those of *Cloacibacterium* sp. 002422665). Bacteroidota CAISCU01 was another bacterium with high expressions of high-affinity *ccoNO* and clade II *nosZ,* which were significantly downregulated under elevated DO concentration. *Paludibacter* (R2_bin.117_r) was the third active member under a microaerophilic condition showing moderate expression for high-affinity oxidase *cydAB* (4.5 times of SCM) and *nrfA* (2.7 times of SCM). In addition, this bacterium had a high expression of *qnorB* gene (8 times of SCM). Thus, this bacterium was capable of utilizing different electron acceptors, particularly

NO, for conserving energy. Increasing DO concentration inhibited the activity of this MAG (**Figs. 5 and 6**).

*Thermomonas* (R1_bin.8_o) utilized high-affinity *ccoNO* genes (expression > 83 times of SCM) as the primary strategy of energy conservation under a microaerophilic condition. Additionally, this bacterium had gene expression of low-affinity oxidases *coxAB* (> 3 times of SCM) (**Fig. 6**). Capability of utilizing both high-affinity and low-affinity terminal oxidases explained the preferential growth of *Thermomonas* bacterium under microaerophilic and alternative redox conditions.

Increasing DO concentration induced the expression of low-affinity oxidases, exclusively for Proteobacteria that possess neither clade II *nosZ* nor DNRA genes (*nrfA* and *octR*). These bacteria also harbored high-affinity oxidases (mainly *ccoNO*) with decreasing expression and other truncated denitrifying genes with low expressions (**Figs. 5** and **6**).

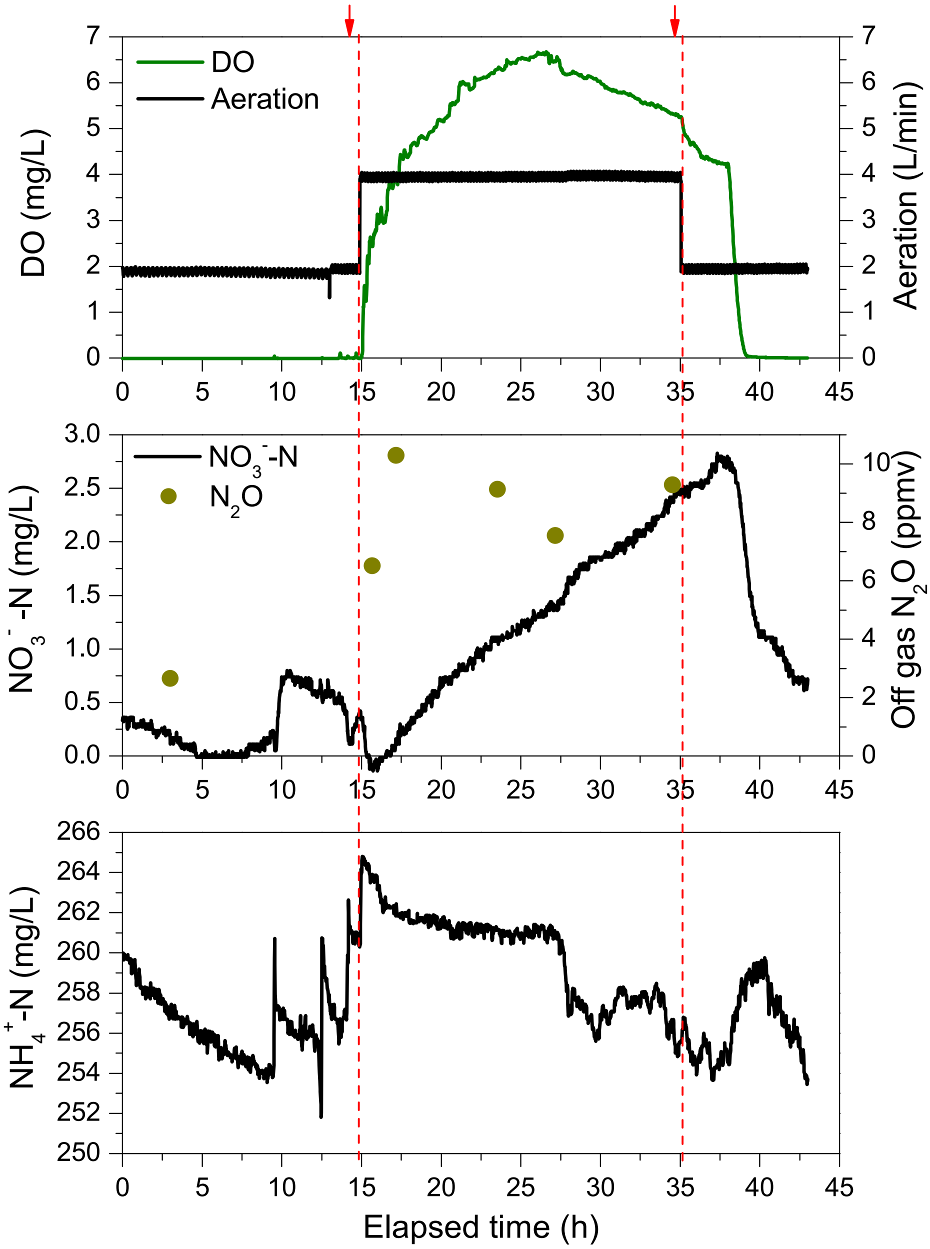


**Fig. S1**: Profiles of aeration rate, DO, $NH_4^+$-N, $NO_3^-$-N, and off-gas $N_2O$ of the MAS system (R2) on days 268 and 269. The red vertical dash lines indicate the time points of doubling and reducing the aeration rates, respectively. The red arrows represent the sampling points for metagenomics (before doubling an aeration rate) and metatranscriptomics (before and after doubling an aeration rate). Aeration volume, DO, $NH_4^+$-N, and $NO_3^-$-N were online recorded while off-gas $N_2O$ was manually sampled and measured by GC-MS.

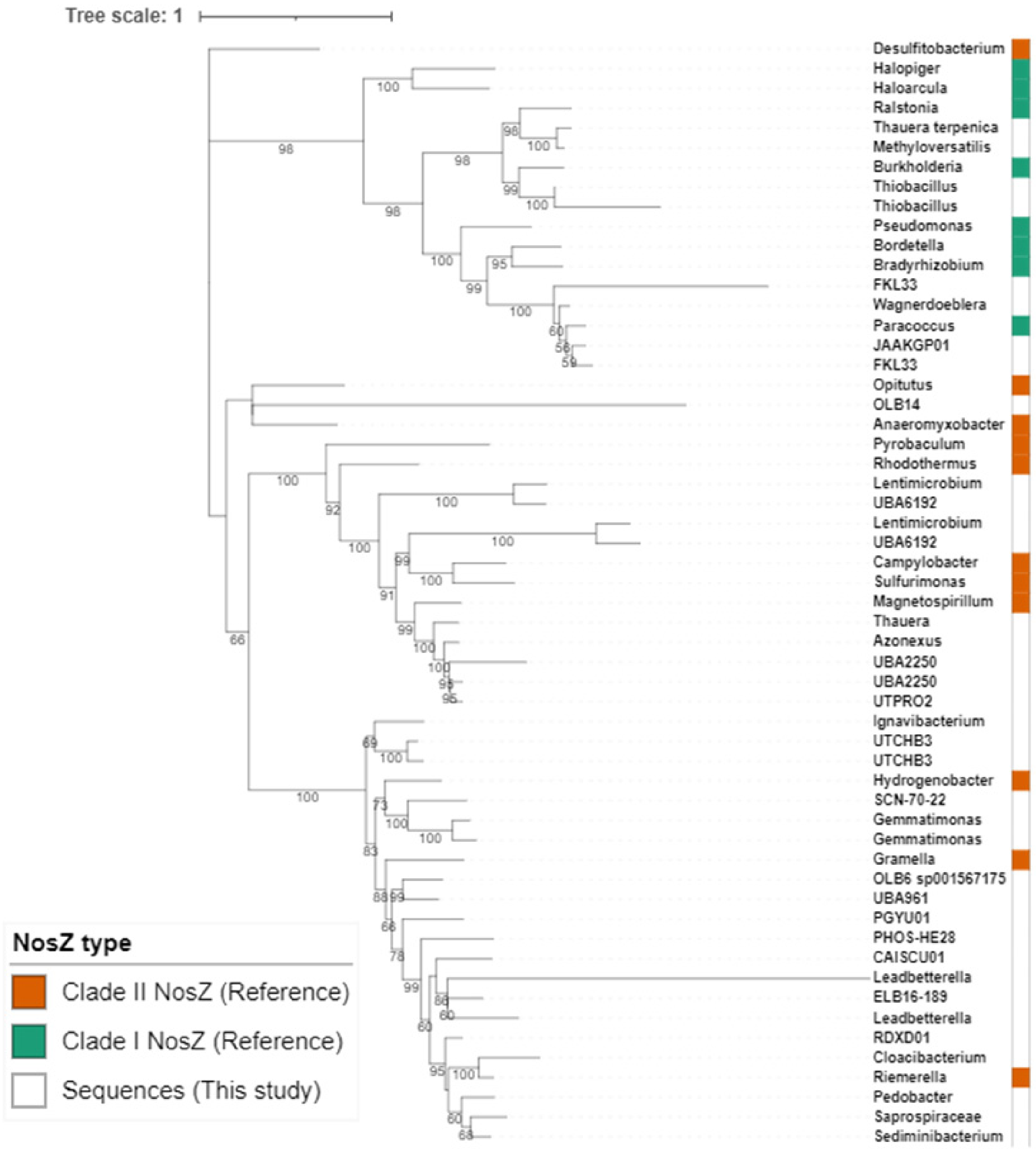


**Fig. S2:** Unrooted phylogenetic tree of NosZ proteins. The tips are labeled with the taxonomy at the genus level. The color strip indicates the references of NosZ types, clade II NosZ (orange) and clade I NosZ (green). NosZ sequences in this study are left blank. Bootstrap values (>50) based on 1000 replications are shown at branch nodes. Reference sequences were selected from a previous study (Sanford et al., 2012) and downloaded from NCBI database (**Table S1**). Sequences were aligned using MUSCLE followed by tree prediction in IQTREE2. The best-fit model, according to BIC, was **Q.pfam+R5**.

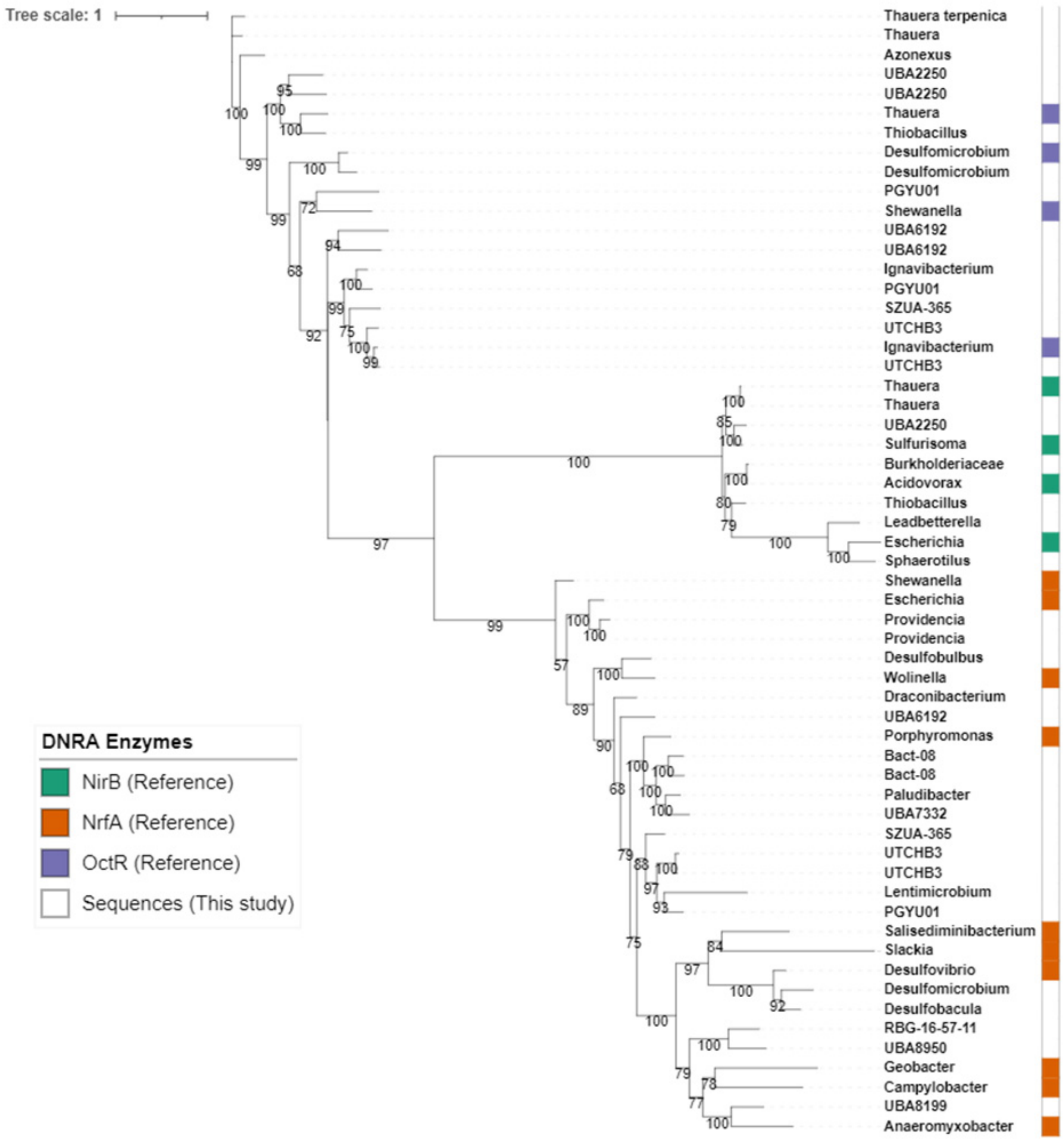


**Fig. S3:** Phylogenetic tree of NrfA, OctR, and NirB proteins (unrooted). The tips are labeled with the taxonomy at the genus level. The color strip indicates the references of NirB (green), NrfA (orange), and OctR (purple). Sequences in this study are left blank. Bootstrap values (>50) based on 1000 replications are shown at branch nodes. Reference sequences for NrfA were selected from a previous study (Welsh et al., 2014). References for NirB and OctR were top Blastp hits downloaded from NCBI. NirB sequence from *E.coli* and OctR sequence from Shewanella are included. Accession numbers of all reference sequences are provided in **Table S1**. Sequences were aligned using MUSCLE followed by tree prediction in IQTREE2. The best-fit model, according to BIC, was **WAG+R5**.

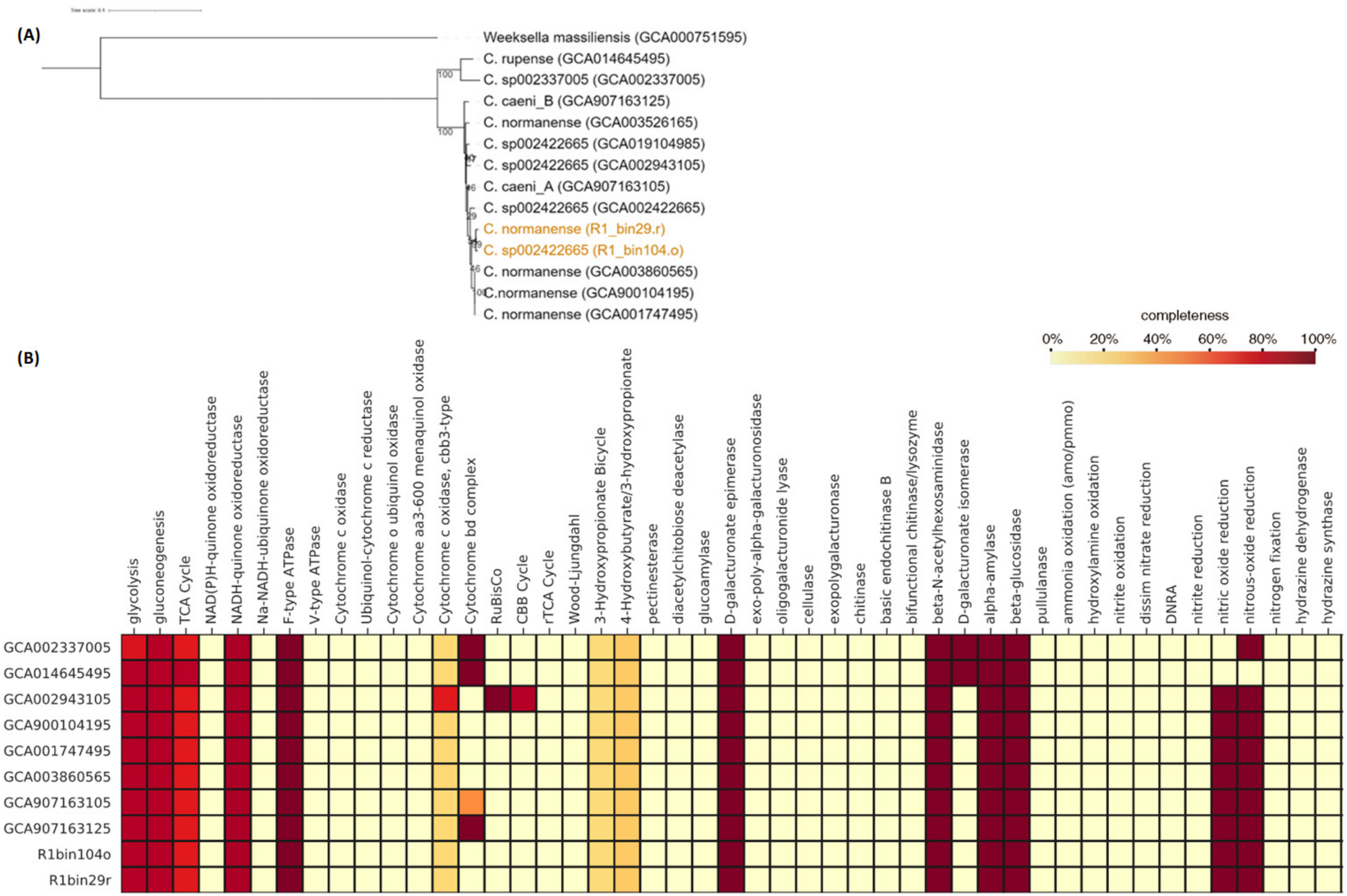


**Fig. S4**: The taxonomy and genotypes of the genus *Cloacibacterium* (Bacteroidota). (A) Phylogenomic tree of all available genomes (completeness > 95% and contamination < 2%) and two MAGs recovered in this study (Orange color). *Weeksella massiliensis* was included as an outgroup. The tips were labeled with taxonomic affiliation classified by GTDB-tk (v2.4.0) and NCBI Genbank assembly accession numbers (in parenthesis). (B) The heatmap shows metabolic pathway completeness calculated by the KEGG Decoder. Each genome was labeled with NCBI GenBank assembly accession number.

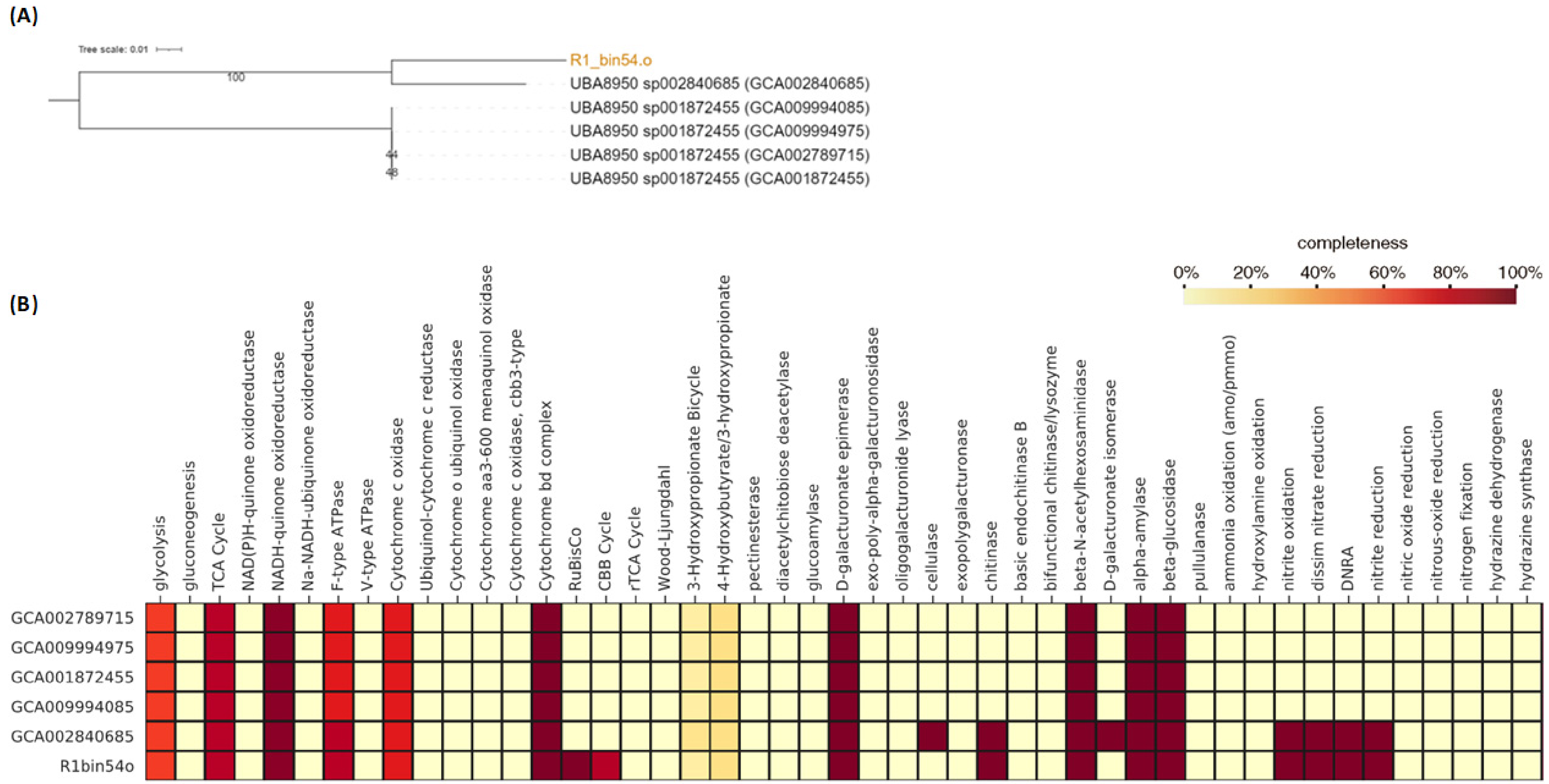


**Fig. S5**: The taxonomy and genotypes of the genus UBA8950 (Chloroflexota). (A) Phylogenetic tree of all available genomes and one MAG recovered in this study (Orange color). The tips were labelled with taxonomic affiliation classified by GTDB-tk (v2.4.0) and NCBI Genbank assembly accession numbers in parenthesis. (B) The heatmap shows metabolic pathway completeness calculated using the KEGG Decoder. Each genome was labelled with NCBI GenBank assembly accession number.

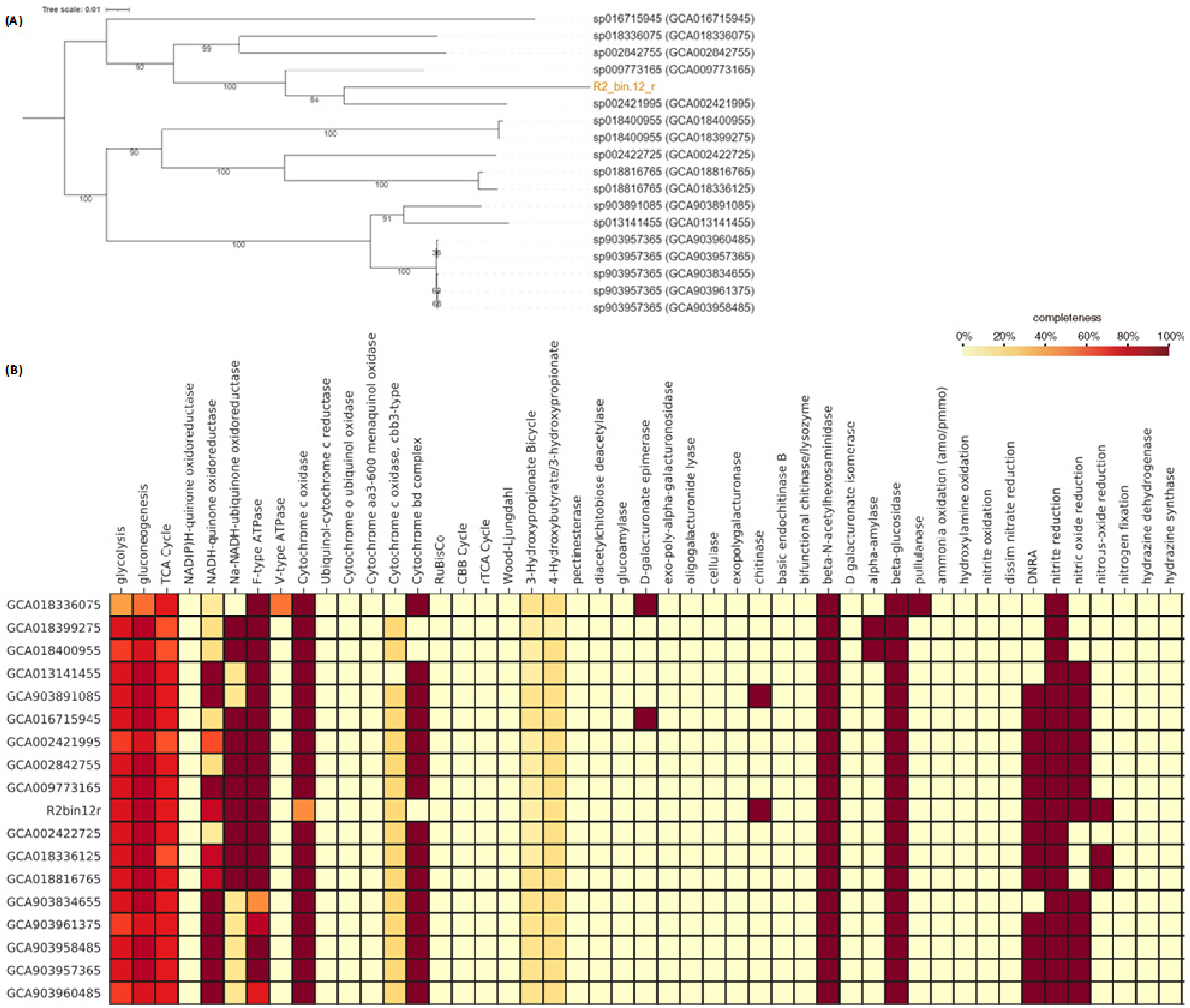


**Fig. S6**: The taxonomy and genotypes of the genus UBA6192 (Bacteroidota). (A) Phylogenetic tree of all available genomes and one MAG recovered in this study (Orange color). The tips were labelled with taxonomic affiliation classified by GTDB-tk (v2.4.0) and NCBI Genbank assembly accession numbers in parenthesis. (B) The heatmap shows metabolic pathway completeness calculated using the KEGG Decoder. Each genome was labelled with NCBI GenBank assembly accession number.

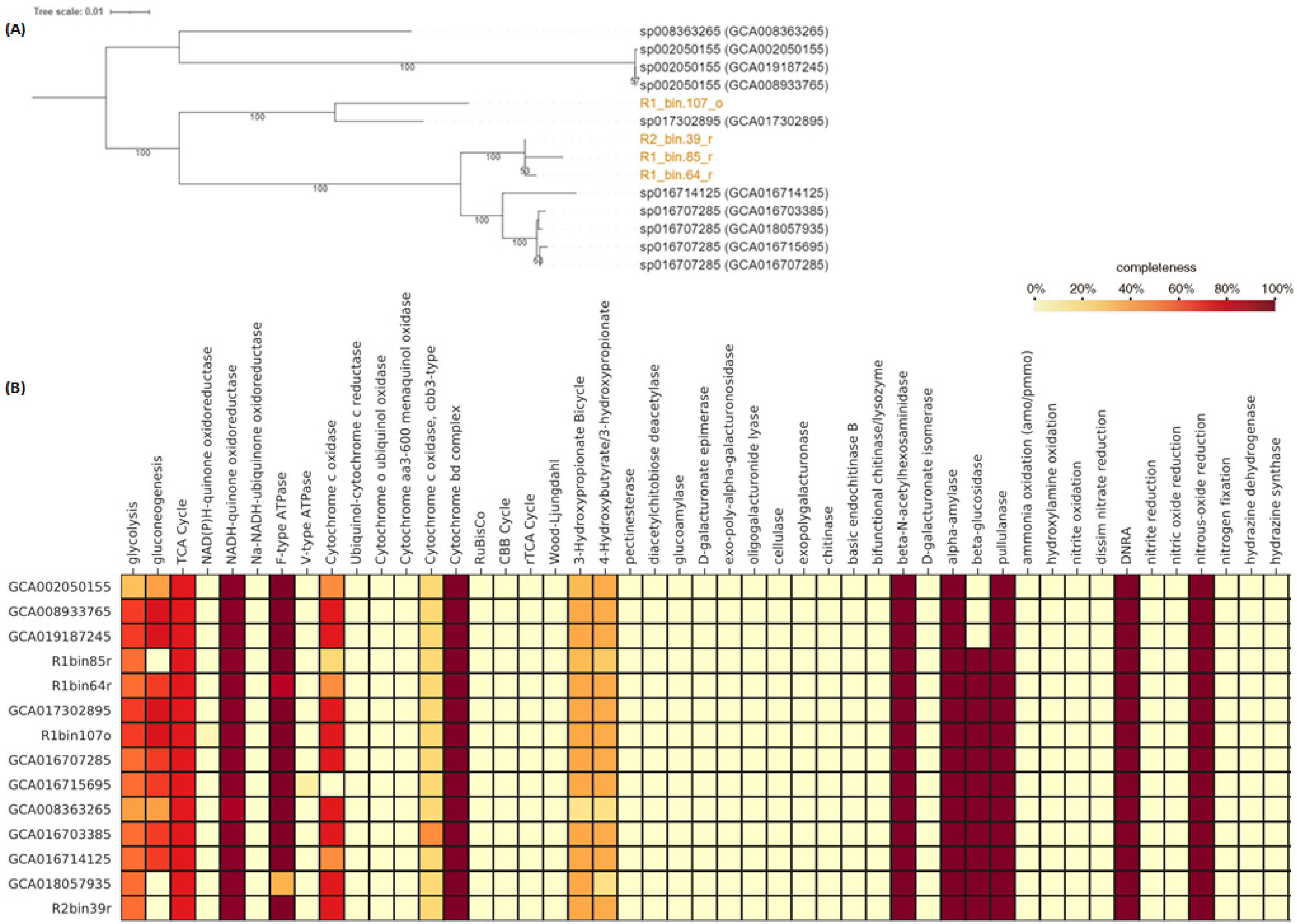


**Fig. S7:** The taxonomy and genotypes of the genus UTCHB3 (Bacteroidota). (A) Phylogenomic tree of all available genomes and four MAGs recovered in this study (Orange color). The tips were labelled with taxonomic affiliation classified by GTDB-tk (v2.4.0) and NCBI Genbank assembly accession numbers in parenthesis. (B) The heatmap shows metabolic pathway completeness calculated using the KEGG Decoder. Each genome was labelled with NCBI GenBank assembly accession number.

**Table S1:** NCBI accession numbers of protein sequences employed to build the phylogenetic trees.

| NosZ references | | | DNRA references | |
|---|---|---|---|---|
| NCBI accession numbers | NosZ types | | NCBI accession numbers | DNRA Types |
| WP_012632836 | Clade II NosZ | | WP_001365050 | NrfA |
| WP_012140537 | Clade II NosZ | | WP_011864650 | NrfA |
| WP_012844504 | Clade II NosZ | | WP_011138866 | NrfA |
| WP_014792165 | Clade II NosZ | | AUR45802 | NrfA |
| WP_004919511 | Clade II NosZ | | WP_004514005 | NrfA |
| WP_011709298 | Clade II NosZ | | ADH98815 | NrfA |
| WP_012962813 | Clade II NosZ | | WP_010937928 | NrfA |
| CAM74903 | Clade II NosZ | | WP_012634152 | NrfA |
| WP_012374624 | Clade II NosZ | | CAG9061977 | NrfA |
| WP_014288022 | Clade II NosZ | | WP_012797970 | NrfA |
| WP_011372928 | Clade II NosZ | | AAN57117 | OctR |
| WP_011222995 | Clade I NosZ | | WP_002931494 | OctR |
| WP_013879286 | Clade I NosZ | | WP_015773582 | OctR |
| WP_012251172 | Clade I NosZ | | MBN8545165 | OctR |
| WP_011083147 | Clade I NosZ | | WP_000049227 | NirB |
| WP_011205031 | Clade I NosZ | | WP_166156952 | NirB |
| WP_011750448 | Clade I NosZ | | WP_121240131 | NirB |
| WP_011914573 | Clade I NosZ | | WP_043741483 | NirB |
| WP_012430154 | Clade I NosZ | | | |

**Table S2:** List of the retrieved MAGs. The version with a higher resolution can be acquired as an excel file.

| No. | Bin.ID | Phylum | Class | Order | Family | Genus | Marker | Number.contig | Completeness | Contamination | Heterogeneity | Genome.size | Longest.contig | N50.contig | Mean.contig | Gene.number | GC.content | Coding.density |
|---|---|---|---|---|---|---|---|---|---|---|---|---|---|---|---|---|---|---|
| 1 | R1_bin.37_o | Thermoplasmatota | Thermoplasmata | Methanomassiliicoccales | Methanomassiliicoccaceae | g_Methanomassiliicoccus | p__Euryarchaeota (UID3) | 680 | 82 | 1 | 0 | 1683672 | 15184 | 2877 | 2476 | 2285 | 1 | 1 |
| 2 | R1_bin.101_r | Gemmatimonadota | Gemmatimonadetes | Gemmatimonadales | Gemmatimonadaceae | g_Gemmatimonas | k__Bacteria (UID2495) | 3 | 99 | 4 | 0 | 4842078 | 4788703 | 4788703 | 1614026 | 4230 | 1 | 1 |
| 3 | R1_bin.104_o | Bacteroidota | Bacteroidia | Flavobacteriales | Weeksellaceae | s_Cloacibacterium sp002422665 | o__Flavobacteriales (UID2815) | 6 | 100 | 0 | 0 | 2804383 | 2786626 | 2786626 | 467397 | 2704 | 0 | 1 |
| 4 | R1_bin.107_o | Bacteroidota | Ignavibacteria | Ignavibacteriales | Ignavibacteriaceae | g_UTCHB3 | k__Bacteria (UID2570) | 17 | 97 | 1 | 0 | 3904923 | 1198518 | 349142 | 229701 | 3547 | 0 | 1 |
| 5 | R1_bin.11_o | Desulfobacterota | Desulfobacteria | Desulfobacterales | Desulfobacteraceae | g_Desulfobacula | c__Deltaproteobacteria (UID3216) | 1481 | 74 | 1 | 20 | 3170540 | 9432 | 2349 | 2141 | 4183 | 0 | 1 |
| 6 | R1_bin.12_r | Riflebacteria | Ozemobacteria | Ozemobacterales | Ozemobacteraceae | g_Rifleibacterium | k__Bacteria (UID2328) | 117 | 96 | 0 | 0 | 5395961 | 249605 | 113119 | 46119 | 4721 | 1 | 1 |
| 7 | R1_bin.13_r | Verrucomicrobiota | Verrucomicrobiae | Pedosphaerales | UBA8199 | f_UBA8199 | k__Bacteria (UID2982) | 166 | 74 | 3 | 0 | 4870708 | 399434 | 106367 | 29342 | 6495 | 1 | 1 |
| 8 | R1_bin.1_o | Patescibacteria | Paceibacteria | UBA9983_A | UBA9973 | g_UBA8515 | k__Bacteria (UID2329) | 90 | 71 | 2 | 92 | 681088 | 28384 | 9897 | 7568 | 746 | 0 | 1 |
| 9 | R1_bin.21_o | Gammaproteobacteria | Gammaproteobacteria | Burkholderiales | Thiobacillaceae | g_PFJX01 | k__Bacteria (UID203) | 336 | 59 | 2 | 0 | 2583953 | 166738 | 49461 | 7690 | 4037 | 1 | 1 |
| 10 | R1_bin.22_o | Desulfobacterota | Desulfobulbia | Desulfobulbales | Desulfobulbaceae | g_Desulfobulbus | c__Deltaproteobacteria (UID3217) | 800 | 82 | 5 | 70 | 2872970 | 17655 | 3953 | 3591 | 3145 | 1 | 1 |
| 11 | R1_bin.24_o | Alphaproteobacteria | Alphaproteobacteria | Rhodobacterales | Rhodobacteraceae | g_Wagnerdoeblera | f__Rhodobacteraceae (UID3340) | 18 | 87 | 0 | 0 | 4859475 | 1114993 | 586703 | 269971 | 5927 | 1 | 1 |
| 12 | R1_bin.25_r | Bacteroidota | Bacteroidia | Chitinophagales | Chitinophagaceae | g_Sediminibacterium | p__Bacteroidetes (UID2591) | 256 | 84 | 0 | 0 | 2663164 | 119326 | 28888 | 10403 | 3100 | 0 | 1 |
| 13 | R1_bin.26_r | Alphaproteobacteria | Alphaproteobacteria | Rhizobiales | Rhizobiaceae | g_FKL33 | o__Rhizobiales (UID3449) | 502 | 72 | 2 | 14 | 4405883 | 188154 | 37631 | 8777 | 5263 | 1 | 1 |
| 14 | R1_bin.27_r | Bacteroidota | Bacteroidia | Flavobacteriales | PHOS-HE28 | g_PHOS-HE28 | k__Bacteria (UID2569) | 300 | 57 | 1 | 0 | 2497145 | 174989 | 29795 | 8324 | 3617 | 1 | 1 |
| 15 | R1_bin.29_r | Bacteroidota | Bacteroidia | Flavobacteriales | Weeksellaceae | s_Cloacibacterium normanense | o__Flavobacteriales (UID2815) | 91 | 100 | 0 | 0 | 2596311 | 343850 | 124245 | 28531 | 2467 | 0 | 1 |
| 16 | R1_bin.31_r | Bacteroidota | Bacteroidia | Bacteroidales | UBA7332 | g_UBA7332 | k__Bacteria (UID2569) | 215 | 90 | 1 | 0 | 3103432 | 225510 | 35825 | 14435 | 3021 | 0 | 1 |
| 17 | R1_bin.32_o | Alphaproteobacteria | Alphaproteobacteria | Rhodobacterales | Rhodobacteraceae | g_JAAKGP01 | f__Rhodobacteraceae (UID3340) | 4 | 98 | 1 | 33 | 4048489 | 4042420 | 4042420 | 1012122 | 4285 | 1 | 1 |
| 18 | R1_bin.33_o | OLB16 | OLB16 | OLB16 | OLB16 | s_OLB16 sp001567115 | k__Bacteria (UID2495) | 1252 | 66 | 0 | 0 | 2884485 | 86410 | 2333 | 2304 | 3766 | 1 | 1 |
| 19 | R1_bin.36_r | Cloacimonadota | Cloacimonadia | Cloacimonadales | Cloacimonadaceae | f_Cloacimonadaceae | k__Bacteria (UID2495) | 41 | 96 | 2 | 0 | 3010179 | 399348 | 256804 | 73419 | 2356 | 0 | 1 |
| 20 | R1_bin.38_r | Patescibacteria | Paceibacteria | UBA9983_A | UBA9973 | g_UBA8515 | k__Bacteria (UID2495) | 9 | 82 | 0 | 0 | 764398 | 452378 | 452378 | 84933 | 777 | 0 | 1 |
| 21 | R1_bin.40_o | Bacteroidota | Bacteroidia | Cytophagales | Spirosomaceae | g_Leadbetterella | o__Cytophagales (UID2936) | 305 | 81 | 4 | 63 | 4665356 | 349168 | 58749 | 15296 | 6326 | 0 | 1 |
| 22 | R1_bin.41_r | Bacteroidota | Ignavibacteria | Ignavibacteriales | Ignavibacteriaceae | g_Ignavibacterium | k__Bacteria (UID2570) | 348 | 65 | 1 | 0 | 2994261 | 93530 | 25055 | 8604 | 3761 | 0 | 1 |
| 23 | R1_bin.42_r | Gammaproteobacteria | Gammaproteobacteria | Burkholderiales | Rhodocyclaceae | s_Azonexus sp012839675 | c__Betaproteobacteria (UID3971) | 100 | 96 | 1 | 43 | 2642200 | 727961 | 264490 | 26422 | 2596 | 1 | 1 |
| 24 | R1_bin.43_o | UBA10199 | UBA10199 | 2-02-FULL-44-16 | JAAZCA01 | f_JAAZCA01 | c__Deltaproteobacteria (UID3216) | 528 | 73 | 2 | 0 | 1792668 | 18051 | 4024 | 3395 | 2003 | 0 | 1 |
| 25 | R1_bin.44_o | Firmicutes | Clostridia_A | Christensenellales | CAG-74 | g_UBA1038 | c__Clostridia (UID1118) | 797 | 61 | 1 | 0 | 1321853 | 6472 | 1665 | 1659 | 1895 | 1 | 1 |
| 26 | R1_bin.45_o | Bacteroidota | Kapabacteria | Kapabacteriales | UBA2268 | g_PGYU01 | k__Bacteria (UID2570) | 61 | 93 | 1 | 100 | 3470364 | 500622 | 166259 | 56891 | 3276 | 0 | 1 |
| 27 | R1_bin.46_r | Bacteroidota | Bacteroidia | Bacteroidales | UBA932 | g_Bact-08 | p__Bacteroidetes (UID2605) | 72 | 91 | 2 | 0 | 2143671 | 244393 | 117369 | 29773 | 2208 | 0 | 1 |
| 28 | R1_bin.48_o | Firmicutes | Bacilli | Lactobacillales | Streptococcaceae | s_Lactococcus petauri | o__Lactobacillales (UID543) | 92 | 100 | 0 | 50 | 2038098 | 203196 | 34104 | 22153 | 2097 | 0 | 1 |
| 29 | R1_bin.49_o | Alphaproteobacteria | Alphaproteobacteria | Rhodobacterales | Rhodobacteraceae | g_Pararhodobacter | f__Rhodobacteraceae (UID3340) | 2 | 100 | 1 | 0 | 4583272 | 4275934 | 4275934 | 2291636 | 4487 | 1 | 1 |
| 30 | R1_bin.50_o | Gammaproteobacteria | Gammaproteobacteria | Burkholderiales | Burkholderiaceae | s_Ottowia sp001897615 | o__Burkholderiales (UID4000) | 20 | 91 | 2 | 50 | 3264912 | 602082 | 271972 | 163246 | 3141 | 1 | 1 |
| 31 | R1_bin.51_r | Gammaproteobacteria | Gammaproteobacteria | Enterobacterales | Enterobacteriaceae | g_Providencia | c__Gammaproteobacteria (UID4442) | 231 | 83 | 0 | 33 | 3824963 | 191367 | 39094 | 16558 | 4130 | 0 | 1 |
| 32 | R1_bin.52_o | Firmicutes | Clostridia | Peptostreptococcales | Acidaminobacteraceae | f_Acidaminobacteraceae | o__Clostridiales (UID1120) | 1 | 99 | 1 | 0 | 3396112 | 3396112 | 3396112 | 3396112 | 3211 | 1 | 1 |
| 33 | R1_bin.54_o | Chloroflexota | Anaerolineae | Anaerolineales | EnvOPS12 | g_UBA8950 | k__Bacteria (UID1452) | 1 | 94 | 4 | 0 | 4743071 | 4743071 | 4743071 | 4743071 | 4318 | 1 | 1 |
| 34 | R1_bin.59_o | Gammaproteobacteria | Gammaproteobacteria | Burkholderiales | Burkholderiaceae | f_Burkholderiaceae | f__Comamonadaceae (UID4119) | 6 | 92 | 0 | 0 | 3295555 | 994485 | 906585 | 549259 | 2956 | 1 | 1 |
| 35 | R1_bin.5_r | Gammaproteobacteria | Gammaproteobacteria | Burkholderiales | Thiobacillaceae | g_Thiobacillus | c__Betaproteobacteria (UID3959) | 142 | 67 | 1 | 0 | 2317804 | 119380 | 44711 | 16323 | 2623 | 1 | 1 |
| 36 | R1_bin.60_r | Gammaproteobacteria | Gammaproteobacteria | Burkholderiales | Rhodocyclaceae | g_UBA2250 | f__Rhodocyclaceae (UID3972) | 22 | 98 | 1 | 40 | 3505927 | 1873752 | 1873752 | 159360 | 3399 | 1 | 1 |
| 37 | R1_bin.62_o | Chloroflexota | Anaerolineae | Anaerolineales | UBA4823 | g_RBG-16-57-11 | k__Bacteria (UID1452) | 920 | 83 | 3 | 0 | 3663385 | 29800 | 5259 | 3982 | 3918 | 1 | 1 |
| 38 | R1_bin.63_r | Firmicutes | UBA994 | JAAYKD01 | JAAYKD01 | f_JAAYKD01 | p__Firmicutes (UID241) | 108 | 84 | 3 | 0 | 2367135 | 356819 | 74449 | 21918 | 2394 | 0 | 1 |
| 39 | R1_bin.64_r | Bacteroidota | Ignavibacteria | Ignavibacteriales | Ignavibacteriaceae | g_UTCHB3 | k__Bacteria (UID2570) | 234 | 93 | 5 | 36 | 4159574 | 174701 | 80127 | 17776 | 4346 | 0 | 1 |
| 40 | R1_bin.65_o | Bacteroidota | Bacteroidia | Chitinophagales | Saprospiraceae | f_Saprospiraceae | p__Bacteroidetes (UID2591) | 55 | 91 | 1 | 50 | 3515869 | 447574 | 207103 | 63925 | 3484 | 0 | 1 |
| 41 | R1_bin.66_r | Synergistota | Synergistia | Synergistales | Aminobacteriaceae | g_Aminivibrio | k__Bacteria (UID209) | 131 | 95 | 3 | 50 | 2683280 | 366389 | 119714 | 20483 | 2555 | 1 | 1 |
| 42 | R1_bin.67_r | Gammaproteobacteria | Gammaproteobacteria | Burkholderiales | Rhodocyclaceae | g_UBA2250 | f__Rhodocyclaceae (UID3972) | 242 | 87 | 2 | 18 | 4030108 | 180367 | 45064 | 16653 | 4145 | 1 | 1 |
| 43 | R1_bin.68_o | Elusimicrobiota | Endomicrobia | Endomicrobiales | Endomicrobiaceae | f_Endomicrobiaceae | k__Bacteria (UID2495) | 4 | 98 | 0 | 0 | 1917760 | 1901614 | 1901614 | 479440 | 1725 | 0 | 1 |
| 44 | R1_bin.69_o | Bacteroidota | Kapabacteria | Kapabacteriales | Kapabacteriaceae | s_OLB6 sp001567175 | k__Bacteria (UID2570) | 828 | 61 | 2 | 25 | 1713765 | 24055 | 2161 | 2070 | 2196 | 0 | 1 |
| 45 | R1_bin.6_o | Gammaproteobacteria | Gammaproteobacteria | Burkholderiales | Burkholderiaceae | g_Rubrivivax | o__Burkholderiales (UID4000) | 14 | 98 | 0 | 100 | 4204232 | 996928 | 912593 | 300302 | 3969 | 1 | 1 |
| 46 | R1_bin.74_o | Bacteroidota | Bacteroidia | Bacteroidales | Lentimicrobiaceae | g_Lentimicrobium | k__Bacteria (UID2569) | 32 | 95 | 3 | 60 | 3714737 | 866350 | 146744 | 116086 | 3362 | 0 | 1 |
| 47 | R1_bin.79_o | Gammaproteobacteria | Gammaproteobacteria | Burkholderiales | Burkholderiaceae | g_SC2-9 | f__Comamonadaceae (UID4119) | 21 | 69 | 0 | 0 | 3294370 | 404554 | 249130 | 156875 | 4183 | 1 | 1 |
| 48 | R1_bin.85_r | Bacteroidota | Ignavibacteria | Ignavibacteriales | Ignavibacteriaceae | g_UTCHB3 | k__Bacteria (UID2570) | 145 | 88 | 3 | 30 | 3616117 | 163018 | 70965 | 24939 | 3840 | 0 | 1 |
| 49 | R1_bin.86_o | Patescibacteria | Dojkabacteria | B142 | o_B142 | o_B142 | k__Bacteria (UID203) | 47 | 71 | 0 | 0 | 981115 | 166061 | 59149 | 20875 | 1317 | 0 | 1 |
| 50 | R1_bin.8_o | Gammaproteobacteria | Gammaproteobacteria | Xanthomonadales | Xanthomonadaceae | g_Thermomonas | f__Xanthomonadaceae (UID4214) | 10 | 99 | 1 | 78 | 2915065 | 1144786 | 665341 | 291507 | 2702 | 1 | 1 |
| 51 | R1_bin.90_r | Desulfobacterota | Desulfovibrionia | Desulfovibrionales | Desulfomicrobiaceae | g_Desulfomicrobium | c__Deltaproteobacteria (UID3217) | 21 | 96 | 1 | 0 | 2620262 | 1970210 | 1970210 | 124774 | 2452 | 1 | 1 |
| 52 | R1_bin.97_o | Gammaproteobacteria | Gammaproteobacteria | Burkholderiales | Rhodocyclaceae | g_UTPRO2 | f__Rhodocyclaceae (UID3972) | 4 | 94 | 1 | 33 | 3270797 | 2666190 | 2666190 | 817699 | 3297 | 1 | 1 |
| 53 | R2_bin.10_o | Bacteroidota | Bacteroidia | Chitinophagales | Chitinophagaceae | g_RDXD01 | p__Bacteroidetes (UID2591) | 1265 | 62 | 0 | 0 | 2628071 | 11891 | 2265 | 2078 | 3277 | 0 | 1 |
| 54 | R2_bin.117_r | Bacteroidota | Bacteroidia | Bacteroidales | Paludibacteraceae | g_Paludibacter | k__Bacteria (UID2569) | 2 | 100 | 1 | 0 | 3389148 | 3382825 | 3382825 | 1694574 | 2683 | 0 | 1 |
| 55 | R2_bin.11_r | Bacteroidota | Bacteroidia | Bacteroidales | UBA932 | g_Bact-08 | p__Bacteroidetes (UID2605) | 73 | 86 | 1 | 0 | 2054625 | 232644 | 65753 | 28146 | 2079 | 0 | 1 |
| 56 | R2_bin.12_r | Bacteroidota | Bacteroidia | Bacteroidales | F082 | g_UBA6192 | k__Bacteria (UID2569) | 11 | 97 | 1 | 50 | 3421419 | 3411058 | 3411058 | 311038 | 2806 | 0 | 1 |
| 57 | R2_bin.13_o | Gammaproteobacteria | Gammaproteobacteria | Burkholderiales | Rhodocyclaceae | g_Methyloversatilis | f__Rhodocyclaceae (UID3972) | 1006 | 80 | 5 | 21 | 2607700 | 17867 | 2892 | 2592 | 3347 | 1 | 1 |
| 58 | R2_bin.15_o | Chloroflexota | Anaerolineae | SBR1031 | A4b | f_A4b | k__Bacteria (UID1452) | 590 | 91 | 4 | 25 | 3751209 | 69781 | 8720 | 6358 | 3737 | 0 | 1 |
| 59 | R2_bin.16_o | Gammaproteobacteria | Gammaproteobacteria | Burkholderiales | Rhodocyclaceae | s_Thauera terpenica | f__Rhodocyclaceae (UID3972) | 10 | 100 | 1 | 0 | 4078798 | 1913548 | 579943 | 407880 | 3773 | 1 | 1 |
| 60 | R2_bin.17_r | Synergistota | Synergistia | Synergistales | Aminobacteriaceae | g_Aminivibrio | k__Bacteria (UID209) | 119 | 97 | 2 | 0 | 2548951 | 273134 | 72573 | 21420 | 2673 | 1 | 1 |
| 61 | R2_bin.18_o | Bacteroidota | Kapabacteria | Kapabacteriales | UBA961 | f_UBA961 | k__Bacteria (UID2570) | 150 | 96 | 1 | 0 | 2910099 | 141496 | 34861 | 19401 | 2403 | 0 | 1 |
| 62 | R2_bin.19_o | Patescibacteria | Gracilibacteria | BD1-5 | UBA6164 | g_UBA7396 | k__Bacteria (UID2329) | 109 | 79 | 1 | 50 | 1249569 | 50762 | 20442 | 11464 | 1219 | 0 | 1 |
| 63 | R2_bin.22_o | Bacteroidota | SZUA-365 | SZUA-365 | SZUA-365 | f_SZUA-365 | k__Bacteria (UID2570) | 1 | 97 | 4 | 0 | 4943508 | 4943508 | 4943508 | 4943508 | 3708 | 1 | 1 |
| 64 | R2_bin.23_o | Bacteroidota | Bacteroidia | Chitinophagales | Chitinophagaceae | g_Sediminibacterium | p__Bacteroidetes (UID2591) | 636 | 75 | 1 | 33 | 2509628 | 59220 | 4435 | 3946 | 2978 | 0 | 1 |
| 65 | R2_bin.25_o | Gammaproteobacteria | Gammaproteobacteria | Burkholderiales | Burkholderiaceae | g_CAISIP01 | o__Burkholderiales (UID4000) | 323 | 65 | 1 | 0 | 3856724 | 71644 | 16167 | 11940 | 3835 | 1 | 1 |
| 66 | R2_bin.28_o | Firmicutes | Clostridia | Peptostreptococcales | Filifactoraceae | s_Acetoanaerobium noterae | o__Clostridiales (UID1120) | 722 | 50 | 0 | 0 | 1198259 | 5860 | 1681 | 1660 | 1706 | 0 | 1 |
| 67 | R2_bin.29_r | Spirochaetota | Spirochaetia | Treponematales | UBA12059 | f_UBA12059 | k__Bacteria (UID2495) | 429 | 92 | 3 | 33 | 2875735 | 1653979 | 1653979 | 6703 | 2825 | 0 | 1 |
| 68 | R2_bin.30_o | Gemmatimonadota | Gemmatimonadetes | Gemmatimonadales | Gemmatimonadaceae | g_SCN-70-22 | k__Bacteria (UID2495) | 23 | 89 | 2 | 0 | 5544337 | 1712291 | 1378532 | 241058 | 5416 | 1 | 1 |
| 69 | R2_bin.31_r | Bacteroidota | Bacteroidia | Cytophagales | Spirosomaceae | g_Leadbetterella | k__Bacteria (UID203) | 120 | 57 | 0 | 0 | 2926753 | 260248 | 87064 | 24390 | 3017 | 0 | 1 |
| 70 | R2_bin.32_o | Gammaproteobacteria | Gammaproteobacteria | Burkholderiales | Burkholderiaceae | s_Sphaerotilus sulfidivorans | o__Burkholderiales (UID4000) | 142 | 94 | 1 | 0 | 3972668 | 160956 | 40522 | 27977 | 3663 | 1 | 1 |
| 71 | R2_bin.35_o | Actinobacteriota | Actinomycetia | Actinomycetales | Microbacteriaceae | s_Leucobacter sp002336855 | o__Actinomycetales (UID1572) | 828 | 73 | 3 | 30 | 1773274 | 9702 | 2273 | 2142 | 2383 | 1 | 1 |
| 72 | R2_bin.36_o | Gammaproteobacteria | Gammaproteobacteria | Enterobacterales | Enterobacteriaceae | g_Providencia | c__Gammaproteobacteria (UID4442) | 177 | 98 | 1 | 40 | 4204146 | 181402 | 42537 | 23752 | 4077 | 0 | 1 |
| 73 | R2_bin.37_o | Gemmatimonadota | Gemmatimonadetes | Gemmatimonadales | Gemmatimonadaceae | g_JAABRT01 | k__Bacteria (UID2495) | 103 | 89 | 2 | 0 | 4512454 | 300935 | 123955 | 43810 | 4265 | 1 | 1 |
| 74 | R2_bin.39_r | Bacteroidota | Ignavibacteria | Ignavibacteriales | Ignavibacteriaceae | g_UTCHB3 | k__Bacteria (UID2570) | 139 | 92 | 4 | 71 | 4026075 | 316685 | 111404 | 28965 | 3504 | 0 | 1 |
| 75 | R2_bin.41_o | Bacteroidota | Bacteroidia | Cytophagales | Cyclobacteriaceae | g_ELB16-189 | o__Cytophagales (UID2936) | 628 | 91 | 1 | 0 | 3603720 | 34836 | 7457 | 5738 | 3700 | 0 | 1 |
| 76 | R2_bin.44_o | Bacteroidota | Bacteroidia | NS11-12g | UKL13-3 | g_CAISCU01 | p__Bacteroidetes (UID2605) | 180 | 100 | 2 | 67 | 3721785 | 105284 | 35428 | 20677 | 3270 | 0 | 1 |
| 77 | R2_bin.47_o | Bacteroidota | Bacteroidia | Bacteroidales | Prolixibacteraceae | g_Draconibacterium | k__Bacteria (UID2569) | 12 | 99 | 3 | 0 | 5718985 | 1971986 | 1725824 | 476582 | 4909 | 0 | 1 |
| 78 | R2_bin.48_o | Spirochaetota | Spirochaetia | Treponematales | UBA12059 | g_SZUA-370 | k__Bacteria (UID2495) | 51 | 92 | 2 | 90 | 3210963 | 431216 | 202242 | 62960 | 3561 | 1 | 1 |
| 79 | R2_bin.4_r | Firmicutes | UBA994 | JAAYKD01 | JAAYKD01 | f_JAAYKD01 | p__Firmicutes (UID241) | 98 | 66 | 1 | 67 | 1637628 | 236943 | 48525 | 16710 | 1841 | 0 | 1 |
| 80 | R2_bin.50_o | Gammaproteobacteria | Gammaproteobacteria | Burkholderiales | Rhodocyclaceae | g_Thauera | f__Rhodocyclaceae (UID3972) | 18 | 92 | 0 | 0 | 4106101 | 1160658 | 447661 | 228117 | 3755 | 1 | 1 |
| 81 | R2_bin.52_o | Bacteroidota | Bacteroidia | Sphingobacteriales | Sphingobacteriaceae | s_Pedobacter sp002399615 | p__Bacteroidetes (UID2605) | 499 | 94 | 0 | 0 | 3257979 | 34510 | 9306 | 6529 | 3253 | 0 | 1 |
| 82 | R2_bin.54_o | Campylobacterota | Campylobacteria | Campylobacterales | Arcobacteraceae | s_Aliarcobacter cryaerophilus_A | k__Bacteria (UID3060) | 30 | 90 | 3 | 56 | 2051938 | 403798 | 164519 | 68398 | 2878 | 0 | 1 |
| 83 | R2_bin.55_o | Gemmatimonadota | Gemmatimonadetes | Gemmatimonadales | Gemmatimonadaceae | g_Gemmatimonas | k__Bacteria (UID2495) | 406 | 85 | 5 | 50 | 3037410 | 34779 | 9620 | 7481 | 2934 | 1 | 1 |
| 84 | R2_bin.61_o | Firmicutes | Clostridia_A | Christensenellales | CAG-74 | g_UBA1038 | o__Clostridiales (UID1120) | 921 | 58 | 0 | 50 | 1680577 | 8408 | 1904 | 1825 | 2284 | 1 | 1 |
| 85 | R2_bin.62_o | Patescibacteria | Paceibacteria | Moranbacterales | UBA1568 | g_SSEF01 | k__Bacteria (UID1452) | 89 | 68 | 0 | 0 | 749941 | 35788 | 12297 | 8426 | 794 | 0 | 1 |
| 86 | R2_bin.64_o | Cloacimonadota | Cloacimonadia | Cloacimonadales | Cloacimonadaceae | g_UBA5553 | k__Bacteria (UID2495) | 520 | 97 | 1 | 100 | 2769591 | 39298 | 8351 | 5326 | 2554 | 0 | 1 |
| 87 | R2_bin.65_o | Alphaproteobacteria | Alphaproteobacteria | Rhizobiales | Rhizobiaceae | g_FKL33 | o__Rhizobiales (UID3449) | 1121 | 78 | 3 | 21 | 4363218 | 246482 | 4283 | 3892 | 5343 | 1 | 1 |
| 88 | R2_bin.66_o | Chloroflexota | Anaerolineae | Anaerolineales | EnvOPS12 | g_OLB14 | k__Bacteria (UID1452) | 43 | 93 | 1 | 0 | 3660944 | 435642 | 180067 | 85138 | 3510 | 1 | 1 |
| 89 | R2_bin.69_o | Bacteroidota | Bacteroidia | Bacteroidales | UBA932 | g_Bact-08 | p__Bacteroidetes (UID2605) | 16 | 96 | 1 | 25 | 2783165 | 804100 | 383399 | 173948 | 2647 | 0 | 1 |
| 90 | R2_bin.6_o | Bdellovibrionota | Bdellovibrionia | Bdellovibrionales | o_Bdellovibrionales | o_Bdellovibrionales | k__Bacteria (UID3187) | 978 | 64 | 2 | 0 | 1893175 | 6938 | 2051 | 1936 | 2638 | 0 | 1 |
| 91 | R2_bin.70_r | Firmicutes | Clostridia | Peptostreptococcales | Anaerovoracaceae | g_UBA1426 | o__Clostridiales (UID1120) | 121 | 98 | 0 | 0 | 2659209 | 390100 | 177977 | 21977 | 2371 | 0 | 1 |
| 92 | R2_bin.71_o | Alphaproteobacteria | Alphaproteobacteria | Rhizobiales | Beijerinckiaceae | g_Bosea | o__Rhizobiales (UID3654) | 913 | 74 | 2 | 30 | 3524863 | 17496 | 4358 | 3861 | 4083 | 1 | 1 |
| 93 | R2_bin.72_o | Gammaproteobacteria | Gammaproteobacteria | Burkholderiales | Thiobacillaceae | g_Thiobacillus | c__Betaproteobacteria (UID3959) | 367 | 95 | 6 | 39 | 2886697 | 58771 | 11763 | 7866 | 3131 | 1 | 1 |
| 94 | R2_bin.74_o | Alphaproteobacteria | Alphaproteobacteria | Rhodobacterales | Rhodobacteraceae | g_UBA1943 | f__Rhodobacteraceae (UID3340) | 138 | 98 | 2 | 29 | 3806111 | 188895 | 49951 | 27581 | 3897 | 1 | 1 |
| 95 | R2_bin.78_o | Spirochaetota | Spirochaetia | Treponematales | Treponemataceae_B | g_UBA12037 | k__Bacteria (UID2495) | 31 | 93 | 0 | 0 | 3586143 | 408076 | 140937 | 115682 | 3797 | 0 | 1 |
| 96 | R2_bin.79_r | Synergistota | Synergistia | Synergistales | Aminobacteriaceae | g_Aminivibrio | k__Bacteria (UID209) | 98 | 95 | 0 | 0 | 2517939 | 268060 | 92590 | 25693 | 2664 | 1 | 1 |
| 97 | R2_bin.82_o | Bacteroidota | Bacteroidia | Bacteroidales | UBA932 | g_Bact-08 | p__Bacteroidetes (UID2605) | 18 | 80 | 0 | 0 | 1951998 | 314314 | 138690 | 108444 | 2051 | 0 | 1 |
| 98 | R2_bin.8_o | Verrucomicrobiota | Verrucomicrobiae | Opitutales | Opitutaceae | f_Opitutaceae | k__Bacteria (UID2982) | 90 | 98 | 1 | 0 | 3084043 | 143535 | 66546 | 34267 | 2796 | 1 | 1 |
